# Dynamics of the cell-free DNA methylome of metastatic prostate cancer during androgen-targeting treatment

**DOI:** 10.1101/2020.04.08.032565

**Authors:** Madonna R. Peter, Misha Bilenky, Ruth Isserlin, Gary D. Bader, Shu Yi Shen, Daniel D. De Carvalho, Aaron R. Hansen, Pingzhao Hu, Neil E. Fleshner, Anthony M. Joshua, Martin Hirst, Bharati Bapat

## Abstract

**Aim:** We examined methylation changes in cell-free DNA (cfDNA) in metastatic castration resistant prostate cancer (mCRPC) during treatment.

**Materials and Methods:** Genome-wide methylation analysis of sequentially collected cfDNA samples derived from mCRPC patients undergoing androgen-targeting therapy was performed.

**Results:** Alterations in methylation states previously implicated in prostate cancer progression were observed and patients that maintained methylation changes throughout therapy tended to have a longer time to clinical progression (TTP). Importantly, we also report that markers associated with a highly aggressive form of the disease, Neuroendocrine-CRPC, were associated with a faster TTP.

**Conclusion:** Our findings highlight the potential of monitoring cfDNA methylome during therapy in mCRPC, which may serve as predictive markers of response to androgen-targeting agents.

## Introduction

Prostate cancer (PCa) is the most common cancer and the second leading cause of cancer-related deaths among males (1, 2). Androgen deprivation therapy (ADT) to reduce systemic androgen levels continues to be the major treatment for aggressive PCa (3). While ADT is initially beneficial, treatment resistance leads to the most lethal form of PCa, metastatic castration resistant prostate cancer (mCRPC) (4). Despite castrate levels of systemic androgens, these tumors can continue to rely on the androgen receptor (AR) pathway to promote tumor growth and metastasis (5). Indeed, androgen-targeting agents, Enzalutamide and Abiraterone acetate, can improve overall and progression-free survival in both the pre- and post-chemotherapy settings (6–9).

Enzalutamide is an antiandrogen that directly inhibits full-length AR. Abiraterone inhibits CYP17A1, which catalyzes extragonadal androgen production (10, 11). Both treatments perform similarly to suppress the androgen pathway, and choice of treatment is often dependent on comorbidities (12). However, primary and eventual secondary resistance to Abiraterone or Enzalutamide remains an ongoing challenge. As a result, the treatment landscape of mCRPC is complex and continues to focus primarily on the order/sequencing of therapies. For instance, recent trials demonstrated early treatment with Abiraterone or Enzalutamide in the hormone-sensitive state could be beneficial (13–15). However, there is a diverse array of molecular drivers contributing to disease heterogeneity amongst mCRPC patients (16, 17), leading to variable response to current treatment strategies. Therefore, reliable biomarkers are needed to facilitate optimal therapy sequences for each patient prior to starting treatments.

Liquid biopsies are emerging as a minimally invasive source of biomarkers, reflecting tumor turnover (18). Numerous studies have explored the potential of circulating tumor cell (CTC) or circulating nucleic acid markers in mCRPC (19). Counting the number of CTCs and/or characterization of molecular alterations (i.e. genomic and transcriptomic) are promising markers that capture tumor heterogeneity (20). For instance, increased CTC number is associated with poor prognosis, and expression of the ligand-independent AR-V7 splice variant by CTCs is associated with resistance to Enzalutamide and Abiraterone (21–23). However, low CTC counts prior to first-line treatment and the need for robust CTC surface markers for their optimal detection remain an ongoing challenge (24, 25). Furthermore, some AR-V7 positive patients may still benefit from androgen therapies, while certain AR-V7 negative patients show variability in treatment response (26, 27). In addition, highly aggressive androgen-independent Neuroendocrine-CRPC (NE-CRPC) tumors express lower levels of AR-V7 than Adenocarcinoma-CRPC patients (28). Therefore, additional markers are needed to identify this subset of patients.

Circulating cell free DNA (cfDNA) can harbor tumor-specific somatic mutations and capture tumor heterogeneity (29). There is a high concordance (>90%) of tumor mutations (i.e. *AR, BRCA2*) between matched cfDNA and mCRPC biopsies (18). Although the proportion of circulating tumor DNA (ctDNA) can vary from 1-2% to ~30% of cfDNA in mCRPC, the sequencing throughput of massively parallel sequencing platforms can sample cfDNA with sufficient coverage (30). For instance, targeted sequencing of *AR* in cfDNA can detect mutations associated with resistance to Enzalutamide or Abiraterone (31, 32). However, genomic aberrations are not the only contributors to molecular and phenotypic heterogeneity in mCRPC. Aberrations in the epigenome are a hallmark of all stages of PCa, including DNA methylation alterations (33, 34). Hypermethylation in promoter regions of several genes, such as *GSTP1*, *APC*, *HOXD3* and *TBX15*, are currently being investigated as diagnostic and prognostic markers in earlier stages of PCa (34, 35). Furthermore, in biopsy tissue from metastatic lesions, distinct DNA methylation patterns were observed between NE-CRPC and Adenocarcinoma-CRPC (28).

CpG methylation can be detected from DNA extracted from tissue and various biofluids (36–38). Presence of promoter methylation in *GSTP1* and *APC* in cfDNA from mCRPC patients is prognostic of overall survival (36, 39). Using an array-based platform, differential cfDNA methylation patterns were observed between Abiraterone responsive versus resistant patients (37). Recently, NE-CRPC tissue-derived methylation signals were shown to be detectable in matched plasma samples (40). While these observations are promising, extensive genome wide analysis of the mCRPC cfDNA methylome during Enzalutamide or Abiraterone treatment has not been performed. To identify circulating DNA methylation changes associated with response to either Enzalutamide or Abiraterone treatment, we monitored sequentially collected cfDNA samples from mCRPC patients, starting from prior to treatment initiation to eventual clinical progression, and performed genome-wide methylome assessment.

Given that biopsy tissue samples are not easily obtainable in the clinical setting, especially bone metastases, we addressed whether analysis of intra-patient cfDNA methylation changes without a priori knowledge of matched tumour methylation patterns, and/or integration with tissue methylation datasets is required to correlate with response to treatment. We identified key genes known to have altered methylation states in PCa and found genomic regions associated with clinical progression. Furthermore, as these changes could be from tumour and non-tumour derived cfDNA, we combined our findings with published data and revealed cfDNA methylation patterns associated with worse clinical prognosis during androgen-targeting treatment.

## Patients and Methods

### Patient cohort

All patients in this study were recruited from the Princess Margaret Cancer Centre Genitourinary Clinic, and informed written consent was obtained in accordance with approved institutional Research Ethics Board protocols from University Health Network (UHN) and Sinai Health System (SHS). The UHN Genitourinary Biobank performed patient recruitment, blood collection and clinical follow-up. All patients were chemotherapy naïve and developed resistance following initial PCa treatments (local therapies and ADT) (**Supplementary Table 1**). A total of 12 Enzalutamide-treated (160 mg daily) and 4 Abiraterone-treated patients (1000 mg daily + Prednisone/Dexamethasone) completed study visits for cfDNA methylome analysis. Blood was collected at three time points: prior to starting treatment (**Visit A**), at 12-weeks during treatment (**Visit B**) and upon clinical progression/treatment change (**Visit C**). While we aimed to collect Visit B around week-12 (+/− 2 weeks), average time from baseline for Visit B was 12.7 weeks (ranging 9 – 17 weeks). Serum PSA levels were obtained throughout treatment, and the lowest PSA level (nadir) was used to determine PSA progression. Clinical progression was determined by several factors, including radiological evidence of additional/worsening metastases and symptomatic changes as assessed by the treating physician.

### Blood sample processing and cfDNA isolation

Blood was collected at each time point in citrate cell preparation tubes (BD Biosciences, San Jose, CA, USA) and samples were processed within 2 hours from collection. Plasma was separated from peripheral blood mononuclear cells (PBMCs) by centrifugation at 1800 × g for 20 minutes. Isolated plasma was divided into 1 mL aliquots and stored at −80°C. Prior to cfDNA isolation, frozen plasma samples were rapidly thawed at 37°C and spun at 16 000 × g for 5 minutes. For each visit, we utilized 4 mL of plasma and cfDNA was isolated using the QIAamp Circulating Nucleic Acid Kit (Qiagen, Hilden, Germany). The concentration of cfDNA was determined using the Qubit dsDNA High Sensitivity Assay Kit (ThermoFisher Scientific, Waltham, MA, USA) and cfDNA fragment distibrution for subset of samples was obtained using the Agilent 2100 Bioanalyzer System. To avoid within-patient batch effects, we isolated plasma cfDNA and performed methylation analysis for all visits from each patient together.

### cfMeDIP-seq protocol

Genome-wide methylation profiling was performed using a published protocol, cell-free methylated DNA immunoprecipitation and high-throughput sequencing (cfMeDIP-seq) (41). Briefly, isolated cfDNA was first end-repaired and A-tailed using the KAPA HyperPrep kit (Roche, Mannheim, Germany), followed by adapter ligation with index adapters (Integrated DNA technologies). Each adapter-ligated cfDNA sample was then spiked with λ phage DNA (ThermoFisher Scientific) as filler DNA, as well as methylated and unmethylated control *Arabidopsis thaliana* (AT) DNA (Diagenode, Denville, NJ, USA) to bring the total DNA amount to 100ng. Prior to immunoprecipitation, 10% of this DNA is saved as input control. The Diagenode MagMeDIP and IPure v2 kits were utilized to enrich and purify methylated cfDNA. Prior to library amplification, qPCR quality checks were performed to ensure: (1) high specificity (>95%) for enrichment of methylated AT DNA over unmethylated AT DNA, and (2) appropriate adapter-ligation. This was followed by adapter-specific amplification of each MeDIP and input control library and gel size selection. All sequencing was performed on the Illumina HiSeq 2500 platform (50-bp single end reads).

### Sequencing data pre-processing and differential methylation analysis

Raw sequencing data was aligned to hg19 reference genome using Burrows-Wheeler alignment (BWA version 0.7.6a, using ‘mem’ option with default settings, except using −M flag). Duplicated reads were marked by Picard tools (https://broadinstitute.github.io/picard/) and collapsed to allow only one copy of the reads from reads that have the same alignment position. Reads mapped to the multiple locations were removed by applying alignment quality threshold (QA>5). As the sequencing data was single-end, the average DNA fragment length was evaluated for both cfMeDIP and input controls using self-correlation technique (42). For each library, the fragment coverage genomic profile was calculated using directional extension of all aligned reads with estimated average fragment length (175bp in our case) using an in-house BAM2WIG custom tool (M. Bilenky, unpublished, see http://www.epigenomes.ca/tools-and-software). Within-patient differential methylation analysis for all comparisons was performed using DMRHunter pipeline developed for cfDNA samples (**Supplemental Methods Section**).

### Statistical analysis of mCRPC-associated DMRs

Spearman correlation analysis was performed using the base package of R (v3.5.3). Receiver operating characteristic (ROC) curve analysis was performed using the pROC package (1.14.0). All data was visualized using the ggplots2 package (v 3.1) and heatmaps from the package ComplexHeatmap (1.2). Pathway analysis is described in **Supplemental Methods Section**.

## Results

### Study design and patient cohort

Tumor methylation signatures can be identified through comparison of cancer patients’ cfDNA with either matched tumor tissue or healthy control cfDNA (41). In the case of mCRPC, biopsy of metastatic lesions in bone is not routinely feasible. Therefore, to identify methylation markers associated with response to therapy, we opted for within/intra-patient differential methylation analysis. We hypothesized that sequential collection of cfDNA from visit A (baseline/pre-treatment) to visit B (week-12) to visit C (clinical progression) would reveal changes in the cfDNA methylome that reflect responses to treatment, with each patient serving as their own internal control. As outlined in **Figure 1**, intra-patient comparison could potentially detect changes in abundance of methylated DNA fragments related to tumor response. For instance, loss of methylated fragments may reflect loss/reduction of tumor cells sensitive to treatment. In contrast, gains in methylated fragments could be associated with gains in treatment resistant tumor cells, since changes in methylome could occur due to non-tumor/treatment related changes. Therefore, we first examined intra-patient differentially methylated regions (**DMRs**) to assess overall changes in cfDNA during treatment. We then integrated these DMRs with tumour-derived methylation datasets.

**Figure 1.**
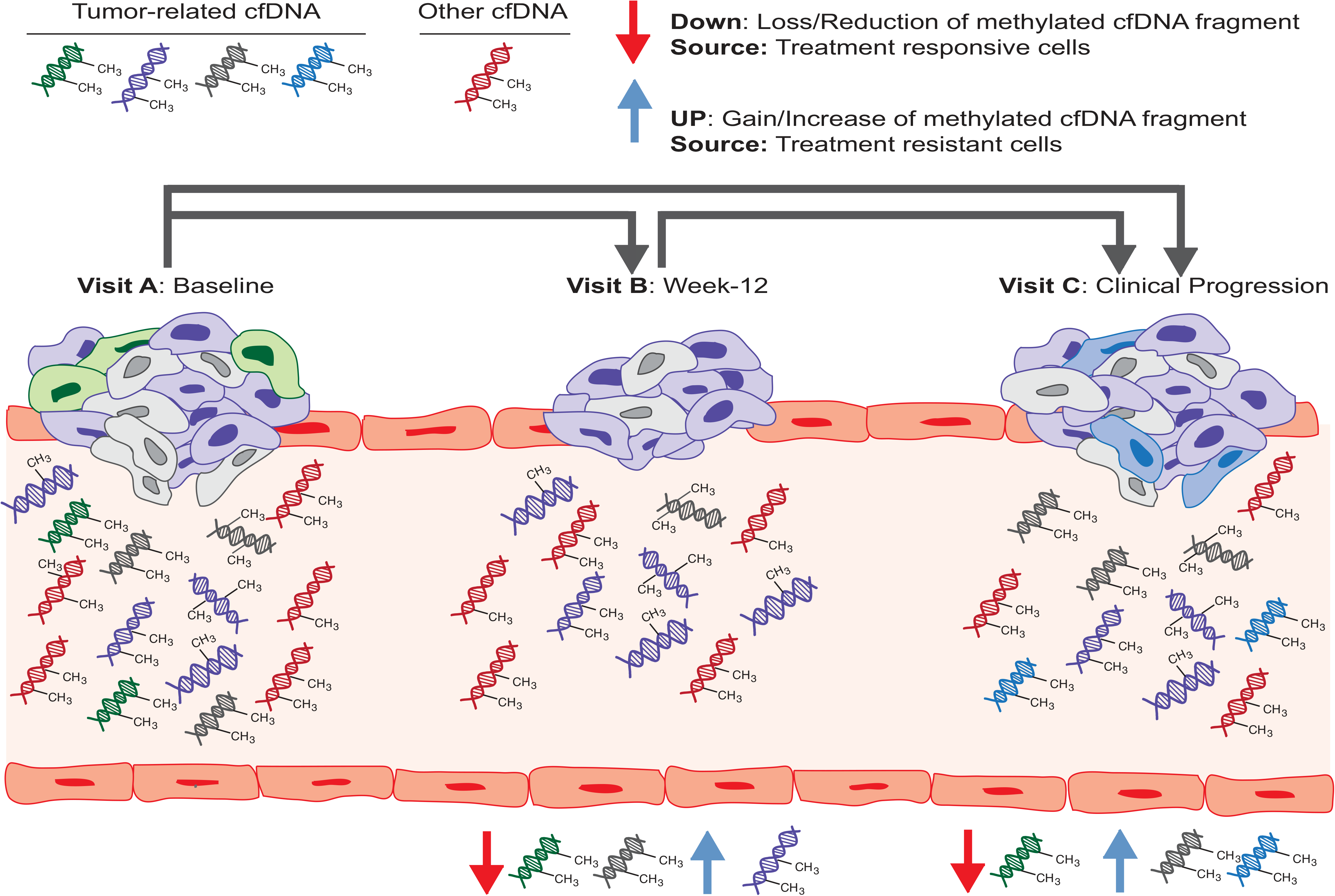
Within patient differential methylation analysis strategy to monitor temporal changes in cfDNA. In order to help identify methylation changes associated with treatment in cfDNA, we opted to perform within-patient methylation analysis to identify differentially methylated regions (DMRs) associated with tumor response and/or resistance to current treatments. Each patient in this study received either Enzalutamide or Abiraterone acetate and blood was collected prior to initiating therapy (**Visit A**), at 12-weeks during treatment (**Visit B**) and upon clinical progression/treatment change (**Visit C**). We applied an established genome-wide method to detect methylated cfDNA fragments followed by extensive analysis to identify DMRs associated with treatment response. We compared all study visits available (Visit B vs A, C vs B, and C vs A) to find losses or gains in methylated cfDNA fragments. For instance, loss of treatment responsive tumor cells (green methylated DNA fragments at visit B) or gains in treatment insensitive/resistant tumor cells (blue DNA fragments at visit C) could be detected with this strategy.

We prospectively collected plasma from mCRPC patients receiving either Enzalutamide (n=12) or Abiraterone acetate (n=4) treatment. In total, 45 blood samples were collected and summarized in **Figure 2a**. Two Enzalutamide-treated patients were unable to provide Visit C samples (P2 and P26) and one Abiraterone-treated patient progressed prior to Visit B (P36). The majority of patients (14/16) presented with bone metastatic lesions upon mCRPC diagnosis, with a subset showing lymph node and soft tissue metastases (**Figure 2b**). Twelve patients demonstrated an initial favorable prostate-specific antigen (PSA) response (≥ 50% decline from baseline, **Figure 2c**), and most patients exhibited maximal PSA response (favorable or plateau) around Visit B (**Supplementary Figure 1**). Overall time to clinical progression (**TTP**) varied for each patient, ranging from rapid/primary resistance to therapy to treatment-driven resistance following initial favorable response (**Figure 2d**).

**Figure 2.**
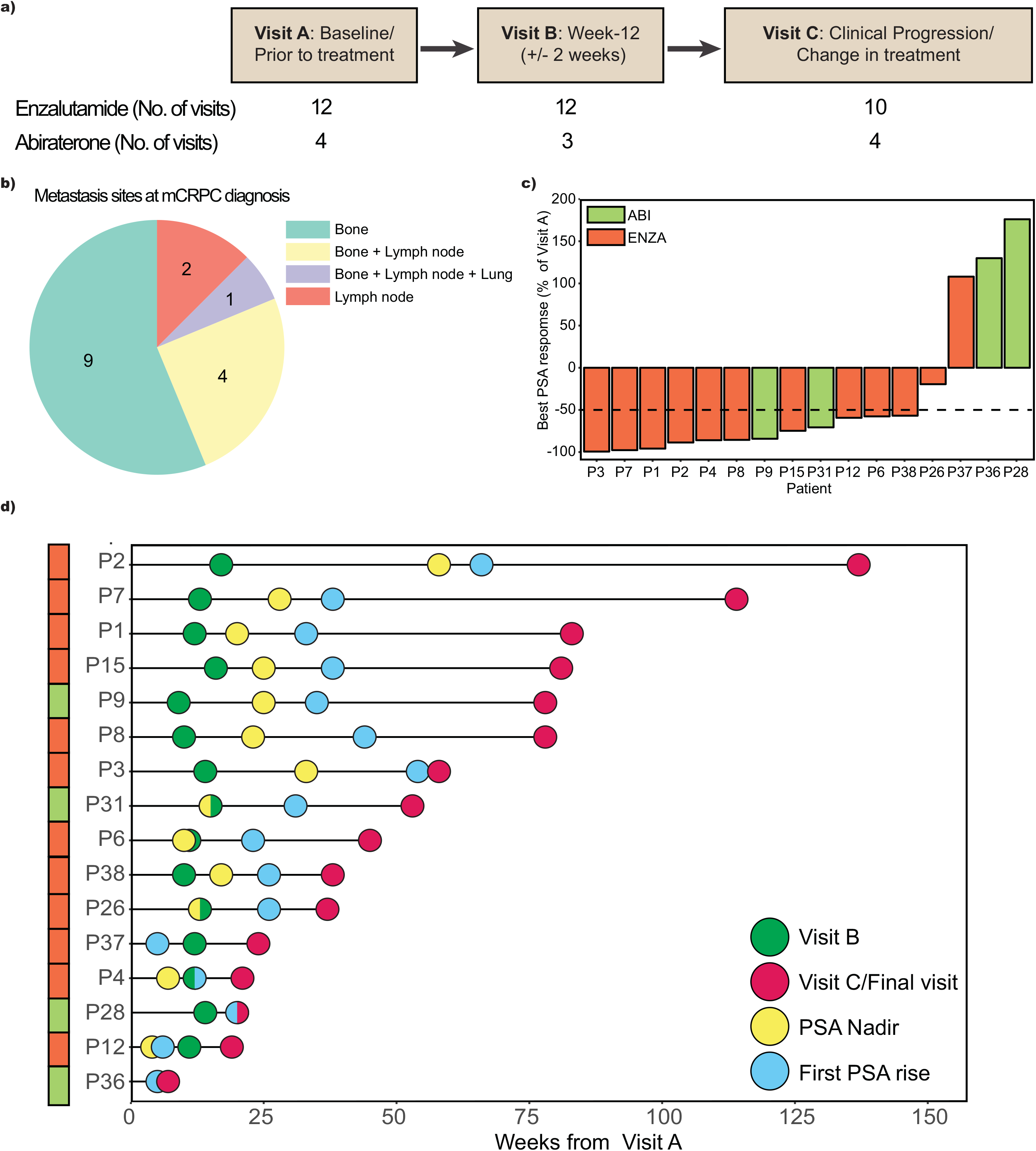
Sample collection and patient clinical follow-up overview. (**a**) Plasma samples from mCRPC patients receiving Enzalutamide (12 patients) or Abiraterone acetate (4 patients) were collected at baseline (Visit A), at week-12 (+/− 2 weeks, Visit B) and upon clinical progression (Visit C). The total number of samples collected at each visit is shown. 2 Enzalutamide-treated patients (P2 and P26) were unable to provide samples for Visit C and 1 Abiraterone-treated patient (P36) progressed prior to Visit B. For these patients, the date of treatment change/clinical progression was recorded for data analysis. (**b**) Pie chart summarizes the distribution of metastasis locations at mCRPC diagnosis. (**c**) Bar plot shows best PSA response for each patient during treatment (nadir), expressed as a percentage of Baseline/Visit A PSA. Dotted line indicates ≥ 50% decline in PSA from Visit A. (**d**) Overall timeline of study follow-up starting from Visit A to Visit C/Final visit is shown for all patients. Green dots represent Visit B, red dots are Visit C or final study visit (if Visit C was not collected), yellow indicates lowest PSA level (nadir) and blue dots show first PSA rise during treatment.

### Performance of cfMeDIP-seq strategy with PCa cell line DNA and cfDNA samples

We utilized an established cfMeDIP-seq approach, which was developed for low amounts of cfDNA and capable of ctDNA fractions as low as 0.001% (41). This technique was able to distinguish methylation patterns between various cancer types, including pancreatic, breast and colorectal cancer. Briefly, methylated cfDNA is enriched using an antibody specific for methylated CpG sites, followed by amplification of adapter-ligated libraries. For each sample, 10% of the DNA was reserved as an input control (without immunoprecipitation).

To benchmark this methodology for PCa samples, we generated a control from genomic DNA derived from PCa cell line, 22Rv1, sheared to the same size as cfDNA (**Supplementary Figure 2**). As an additional control, we spiked all samples with methylated and unmethylated *Arabidopsis thaliana* (AT) DNA, and confirmed increased recovery of methylated AT DNA (~80%) compared to unmethylated DNA (<0.06%) (**Supplementary Figure 2a**). Spiked AT DNA was added following sequencing adapter ligation to cfDNA and not detectable during sequencing. Increased enrichment of known methylated genes/regions in the genome of 22Rv1 cells, (43) including the *CDKN2A* gene body and the first exon of *TGFB2,* compared to the unmethylated promoter of *HOXD8* was confirmed (**Supplementary Figure 2b**).

We performed cfMeDIP-seq on all 45 samples collected. The average starting amount of dsDNA was 27 ng, and ranged from 9 to 50 ng, with no significant differences in cfDNA amounts across visits and treatments (**Supplementary Figure 2c**). Following alignment and base quality filtering, 48-62% of the cfMeDIP reads and 80-85% of input control reads remained for DMR analysis (**Supplementary Figure 2d-e**). A majority of cfMeDIP-seq sequence reads aligned within gene bodies (major transcript starting from transcriptional start site/TSS to transcription termination site/TTS) and intergenic regions, as well as CpG Island (CGI), CGI shores, and TSS/promoter regions (TSS +/−1.5Kb), together representing the entire genome (**Supplementary Figure 2f**). As expected, enrichment of methylated cfDNA in the MeDIP fraction over input control fraction was observed, especially within CpG rich regions. Similar distribution of regions was observed between Enzalutamide and Abiraterone treated patients (**Supplementary Figure 2 g-h**).

### Intra-patient differential methylation analysis

To identify treatment-related differentially methylated regions (DMRs), we developed a DMR calling strategy for intra-patient analysis termed “cfDNA DMRHunter” (See **Supplemental methods** and **Supplementary Figure 3**). As location-specific cfDNA recovery sensitivity is unknown *a priori*, we used input control libraries to ensure uniform cfDNA read coverage across the whole genome. DMRHunter also uses the input controls to remove alignment artifacts from analysis. After filtering low quality and PCR duplicated reads, DMRHunter builds a background model and identifies differentially methylated CpGs (DMCs) across the genome. Adjacent DMCs with the same differential methylation state (UP or DOWN) were merged to identify DMRs with stringent z-score (>1.5), average read count (≥10) and false discovery rate (FDR = 0.01) cut-offs. There were a large number of DMRs across all comparisons and treatments, ranging from 93 000 to 154 000 DMRs (**Figure 3a**) with no significant difference in median number of DMRs between comparisons and treatment (**Figure 3b-c**). The overall median DMR size was 74 bp, ranging from 2bp (single CpG sites) up to 4163 bp.

**Figure 3.**
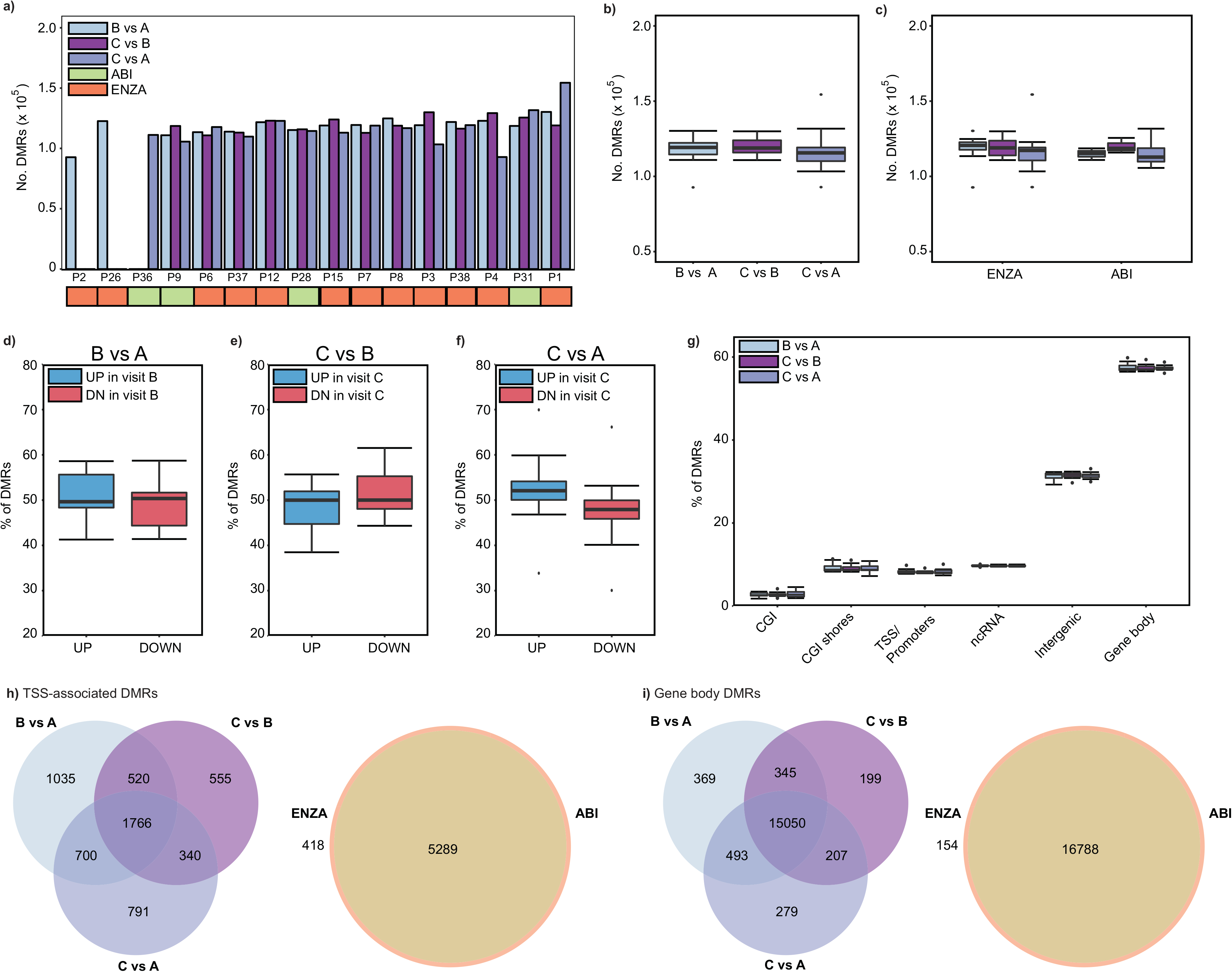
Summary of intra-patient DMR analysis. (**a**) The total number of DMRs for all patients, visit comparison and treatment. (**b**) Boxplot summarizing median, first and third quartile of all comparison DMRs. Whiskers represent the highest and lowest number of DMRs. (**c**) DMR totals were further stratified by treatment type. Boxplots illustrate percentage of DMRs that increased or decreased in methylation in (**d**) B vs A, (**e**) C vs B and (**f**) C vs A. (**g**) The overall proportion of DMRs that were identified near/within CGI, shore, TSS/promoter, ncRNA, intergenic and gene body regions is shown for all visit comparisons (In order of B vs A, C vs B and C vs A). (**h**) Venn diagrams showing the overlap of protein coding genes with DMRs near their promoters is shown for all visit comparisons and between Enzalutamide- and Abiraterone-treated patients. (**i**) Similarly, the overlap of protein coding regions with DMRs within gene bodies is shown.

We next examined the directionality of these changes across comparisons. Conventionally, hypermethylation or hypomethylation is the typical terminology for methylation changes. However, for cfDNA, it is unclear whether these changes reflect a gain/loss of methylated CpGs at these specific genomic sites or changes in DNA fragment abundance, reflecting an altered cellular abundance contributing to the cfDNA pool. For our analysis we opted to define these DMRs as gains (UP) or loss (DOWN) in methylated fragments. There was no significant difference in the overall proportion of regions with increased or decreased methylation in all visit comparisons (**Figures 3d-f**). The majority of DMRs were found within gene bodies (protein coding and pseudogenes) and intergenic regions (**Figure 3g**). DMRs near/within transcription start sites (TSS)/promoters and CpG Islands (CGI), as well as non-coding RNA (ncRNA), were found, but with lower abundance. We chose to focus on DMRs within promoter and gene body regions, as CpG methylation changes in these locations are known to be associated with gene expression changes (44).

### Methylation changes within promoter and gene body regions

The majority of DMRs within promoters (TSS +/− 1.5 Kb) were associated with protein coding (PC) genes, followed by long non-coding RNA (lncRNA), pseudogenes (Pseudo) and short non-coding RNA (sncRNA) regions (**Supplementary Figure 4a**). We observed minimal overlap among promoters of different gene/ncRNA categories, and overall distribution of DMRs in these regions did not vary between patients and visits (**Supplementary Figure 4b**). We focused on protein coding genes with at least one promoter DMR. Analyzing recurrence of these genes, we found that many of these genes were found in less than 5 patients for each comparison (**Supplementary Figure 4c-e**). We examined the most common protein coding genes (with DMRs in ≥ 5 patients) and found many genes overlapping across visit comparisons, with the majority shared between Abiraterone and Enzalutamide groups (**Figure 3h**). The most common genes showing either increases or decreases in promoter methylation are in **Supplementary Figure 4f**, with certain genes implicated in PCa or other cancers. For instance, pyrroline-5-carboxylate reductase 1 (*PYCR1*), which is involved in proline biosynthesis and can promote tumor cell growth (45), often demonstrated gain of methylation at visit C. Pathway enrichment analysis of promoter DMRs (**Supplementary Figures 5 and 6**) showed cancer-related pathways such as Wnt and PI3K signaling were observed, with neuronal pathways being the most frequent. Several promoter associated DMRs were only found among Enzalutamide-treated patients (**Figure 3h and Supplementary table 2)**. The most frequently differentially methylated promoter region (6-7 patients, 20 comparisons) within the Enzalutamide cohort was *FGFR1*, which is overexpressed in PCa and implicated in metastasis (46).

We also examined the distribution of DMRs within gene bodies/known transcripts. The majority of these DMRs were associated with protein coding gene bodies across (**Supplementary Figure 7a-b**). In contrast to promoter DMRs, many genes with body DMRs were shared across patients (**Supplementary Figure 7c-e**). Similarly, we focused on commonly altered gene bodies and found extensive overlap between visit comparisons and treatments (**Figure 3i**). We further examined net methylation change across these genes by scoring the ratio of increased to decreased methylation within each gene body, excluding genes that had no net change (i.e. equal number of UP and DOWN trends). We found 91 genes that were commonly altered across all comparisons (**Supplementary Figure 7f**). In most cases, regardless of visit comparison, the methylation pattern for these genes clustered closely for the same patients. One such group of genes with increases in methylation at Visit C included *NFATC1*, which can promote tumorigenesis (47), and a histone deacetylase, *HDAC4*. While several Enzalutamide-specific genes were found (**Supplementary table 3**), common pathways including several differentiation-related and neuronal pathways were found among gene-body associated DMRs (**Supplementary Figures 8 & 9**).

Overall, examining the most frequently altered promoters or gene bodies revealed a few cancer-related genes, but changes across all visit comparisons were variable. We assessed the overall promoter methylation status of genes that have been established as differentially methylated in PCa (**Figure 4a**), including *GSTP1*, *TBX15*, *AOX1* and members of the HOX family of transcription factors (34). Among the most common differentially methylated gene promoters were genes involved in tumorigenic processes, such as *RUNX3*, *RGS12* and *FBP1*, with several differentially expressed in NE-CRPC (28), including the neuronal transcription factor *PHOX2A* (48) and a regulator of apoptosis, *CTBP2* (49). DMRs in genes associated with diagnosis and prognosis could reflect tumor cell response to treatments. While differential promoter methylation events in the well-established PCa-specific hypermethylated gene, *GSTP1*, were infrequent (**Figure 4b**), decreased gene body methylation coinciding with treatment response (i.e. PSA reduction) was shown in some patients (i.e. P3). Similar methylation regions were also observed in the PCa cell line LNCaP (50). Similarly, changes in *TBX15* methylation (1^st^ intron) were associated with response to treatment (i.e. PSA) in certain patients (**Figure 4c**). While promising, there were no consistent DMRs in these genes that could classify patients by androgen-targeting treatment response. To further refine potential treatment associated methylation changes, locations within the genome that experienced fluctuations in methylation levels across all visits were next explored.

**Figure 4.**
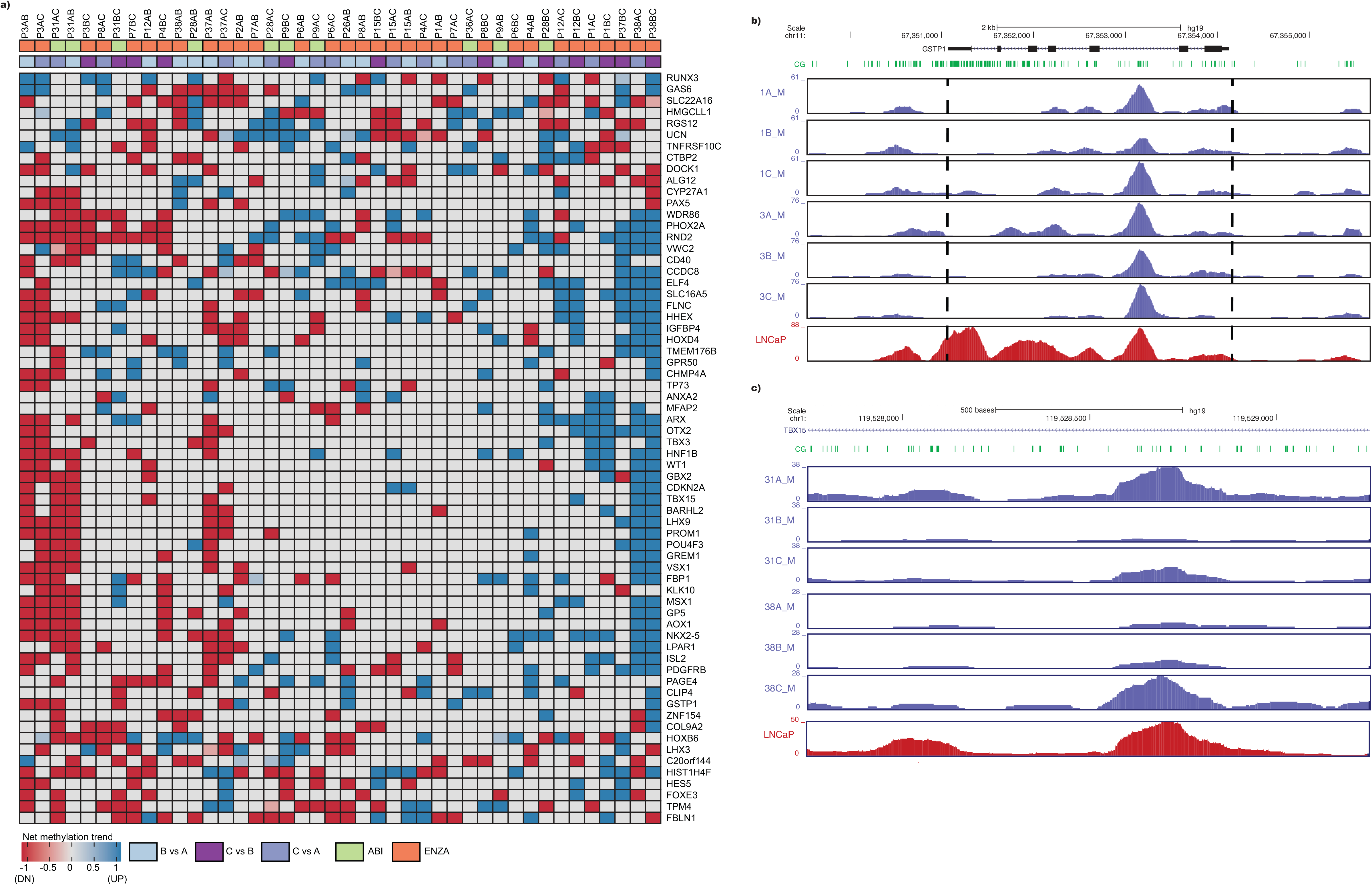
Key prostate cancer related methylation changes observed in cfDNA. (**a**) Heatmap shows methylation trends in promoter regions for all patients and comparisons of the most frequently methylated genes in PCa. For each gene, the proportion of DMRs with increased (blue) and decreased (red) methylation was calculated and the net change is shown. Grey indicates no net change in methylation. Representative peaks show methylation levels (normalized read counts) of (**b**) *GSTP1* (black dotted lines indicating gene body) and (**c**) *TBX15*. LNCaP cell line MeDIP-seq data is also shown.

### Common CpG sites with differential methylation across treatment visits

To find potential “treatment sensitive” CpG sites in the genome, we next examined differentially methylated CpG sites (DMCs) among patients that completed all study visits, specifically CpG sites that fluctuated in ≥2 visit comparisons. DMCs were analyzed as DMR size for similar genomic regions varied within and across patients. We stratified the proportion of DMCs that were uniquely found in single visit comparisons and those shared across all visits (Scenarios 1-6 in **Figure 5a**). We identified DMCs present in two or more visit comparisons for each patient (Scenarios 7-14) and associated them with gene promoters and bodies. Analysis of gene promoters containing DMCs revealed minimal overlap amongst patients, with P4, P31, P38 and P3 showing the highest overlap of 8 shared genes: *NPBWR1*, *ZSCAN12*, *PCDHGA11*, *PHOX2A*, *TBX10*, *TEX28*, *TKTL1* and *TSPAN32* (**Supplementary Figure 10a**). In contrast, we observed a high degree of overlap amongst patients when looking at DMCs within gene bodies, with the exception of P3 (**Supplementary Figure 10b**). Interestingly, we also noted a positive correlation between the proportion of DMCs shared across all visits and TTP (**Figure 5a**).

**Figure 5.**
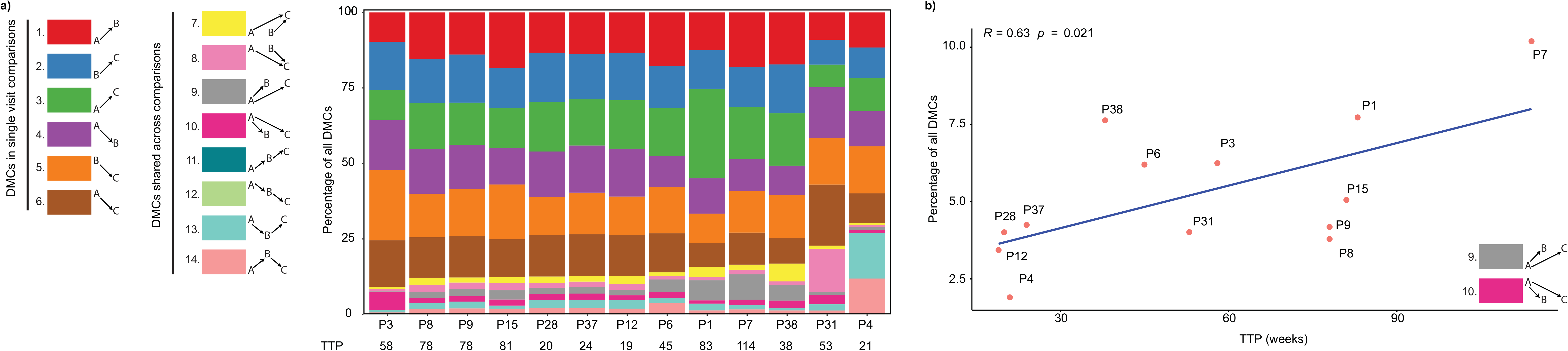
Common differentially methylated CpGs (DMCs). (**a**) For patients that were able to provide samples for all three visits, the proportion of CpG sites that were differentially methylated in a single comparison and shared across 2 or more comparisons is shown. 14 possible scenarios are represented by different colors. TTP in weeks are also indicated for each patient. (**b**) Spearman correlation analysis of the proportion of DMCs that were found to increase or decrease in B/C vs A (scenarios 9 & 10) against time to progression (Spearman rho and p values are shown).

We examined 3 major combined scenarios observed among 14 total: (1) increase/decrease in methylation in visit C vs B and A (scenarios 7 and 8) (**Figure 5a**); (2) increase/decrease in B and C vs A (scenarios 9 and 10,); (3) those that increase /decrease in B vs A and then return to same levels in C (13 &14). The proportion of these three combined scenarios was correlated with time to progression (TTP), and we found that those patients with more DMCs that increased/decreased at visit B and maintained at visit C (scenarios 9 and 10) had a longer TTP (**Figure 5b**). Although scenarios 7 & 8 did not correlate with TTP, those patients with a higher proportion of scenarios 13 & 14 had a possible trend towards faster TTP (**Supplementary figure 10 c-d**, p=0.05). This suggests that genome-wide cfDNA methylation dynamics can reflect response to treatment in mCRPC patients. Importantly, changes in methylation levels of these specific regions/genes could potentially be used as a monitoring tool during androgen-target treatment.

### Neuroendocrine-CRPC related cfDNA DMRs

Thus far, fluctuations in DMRs were analyzed for all cfDNA signals, which could include tumour and non-tumour derived DNA. As many of the DMRs were associated with genes implicated in several neuronal pathways and differentiation, we integrated cfDNA DMRs with those identified from biopsy samples from NE-CRPC versus Adenocarcinoma-CRPC in an independent mCRPC patient series (28). While this published dataset utilized reduced representation bisulfite sequencing (RRBS), which primarily covers CpG rich regions (i.e. CGIs), several of these DMCs were found within cfDNA DMRs (29 355 CpG sites out of 84 930 in NE-CRPC). In order to assess whether changes in these regions at earlier time points are associated with overall response to androgen-targeting agents, we examined the overlap between DMRs from the A vs B comparison and NE-associated DMRs. As P36 progressed prior to visit B (**Figure 2d**), we included this patient in the analysis (examining A vs C). In this case, we examined the changes in methylation from the perspective of visit A. That is, increased methylation in visit A vs B/C (UP in A) or decreased methylation in visit A (DOWN in A). We proposed that the abundance of methylated cfDNA fragments from certain regions may reflect key genes/regions to monitor prior to initiating therapy. The overall proportion of these DMRs that were increased or decreased in methylation at A vs B (or C for ID 36) varied across the patients (**Figure 6a**). Interestingly, we noted that these DMRs (A vs B) appeared to be related to TTP. We examined the ratio of DMRs containing NE-CRPC associated CpG sites with more methylated fragments in Visit A to those with less methylation. In order to further delineate key regions, we applied various FDR thresholds to these DMRs to examine the effect on correlation with TTP (**Figure 6b**). We found an optimal set of regions that were negatively correlated with TTP: that is, the more methylated cfDNA fragments containing these NE-CRPC associated CpGs, the faster/shorter the TTP (**Figure 6c**). In a follow-up study analyzing matched biopsy and plasma methylation signals from 5 Adenocarcinoma-CRPC and 6 NE-CRPC patients, several prostate cancer and Neuroendocrine related methylation regions were found in cfDNA (40). We also integrated our A vs B DMRs with this dataset and found similar correlation with TTP (Spearman R = −0.56, p = 0.024). To determine whether the amount of NE-CRPC related methylation patterns at visit A was associated with a faster TTP, we performed ROC analysis for various TTP cut-offs (ranging from ≤20 to 35 weeks). We found that an increased ratio of NE-CRPC DMRs UP in visit A can stratify patients that progressed as early as 25-weeks (area under the curve, AUC: 0.927, CI: 0.773-1) (**Figure 6d**). These findings confirm the detection of NE-CRPC related methylation signatures in cfDNA, but also show for the first time that these sites may serve as potential predictive markers of resistance to androgen-targeting agents.

**Figure 6.**
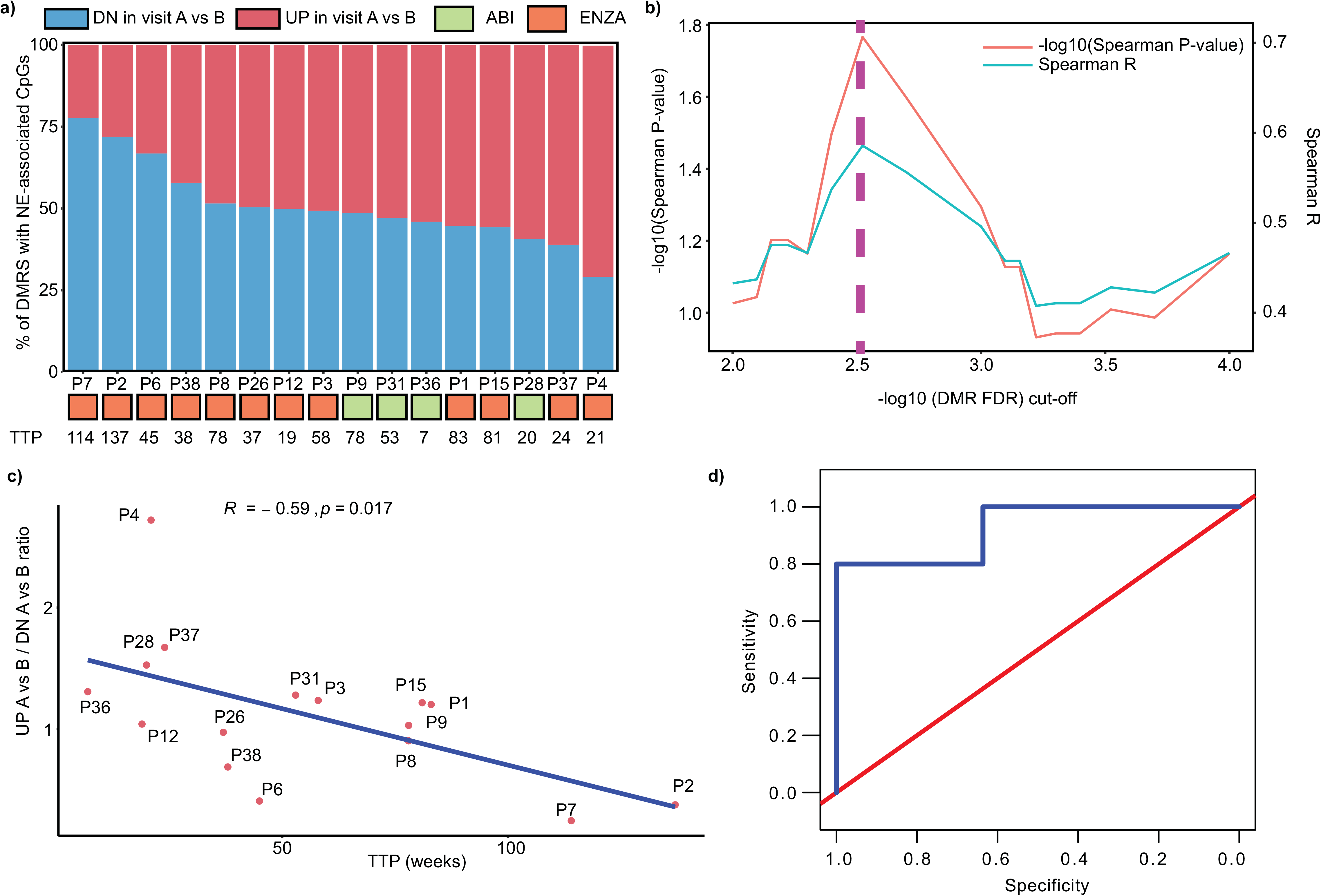
NE-CRPC related DMRs in cfDNA. (**a**) For each patient, the proportion of DMRs that demonstrated methylation differences in NE-CRPC associated regions in visit A vs B. (**b**) For DMRs that contained NE-CRPC associated CpGs, various FDR thresholds were applied to assess for optimal regions that correlated with TTP. (**c**) Using the optimal FDR cut-off, the ratio of NE-associated DMRs with increased methylated fragments in visit A vs B to decreased fragments was correlated with TTP (Spearman rho and p value is shown). For P36, A vs C comparison was used as this patient progressed prior to visit B. (d) ROC analysis shows optimal AUC (0.927, 95% CI: 0.773-1) that stratifies patients according to TTP (<25 weeks).

## Discussion

In this study, we showed that methylome analysis of the cfDNA from mCRPC patients can capture treatment-related epigenetic alterations, with and without integration with tissue derived methylation data. We found changes in well-established PCa methylation markers associated with response to treatment. Importantly, we demonstrated that cfDNA contains NE-CRPC related methylation signals, which have the potential to be utilized as predictive markers of treatment response. Methylation of several oncogenic driving pathways were detected in cfDNA across time points, suggesting that methylome analysis of patients’ liquid biopsies can serve as a monitoring tool during treatment, as well as highlight potential targets for future therapies.

A key advantage of liquid biopsy-based versus tissue-biopsy approaches is the ability to capture tumor heterogeneity. To date, much focus has been placed on genomic alterations, which can help identify *AR* mutations related to resistance to Abiraterone/Enzalutamide (31, 32). However, few studies have highlighted that cfDNA methylation markers may help further stratify patients (36, 37). In a recent study of 600 mCRPC patients receiving docetaxel, increased methylated *GSTP1* at baseline was associated with longer overall survival, and loss of methylation after two cycles of treatment corresponded with longer time to PSA progression (51). Furthermore, patients that exhibited loss in methylated *GSTP1* after two cycles of treatment had better overall survival. In this study, we observed some cases in which there were losses of *GSTP1* methylation in certain treatment responsive patients (PSA reduction). We also examined other tissue-based methylation markers (promoter and gene body), including *TBX15* and *RUNX3* (34), and similar heterogeneity amongst patients was observed. This in part could be explained by differing mechanisms between docetaxel and androgen-targeting agents. Due to intra- and inter-tumor heterogeneity (52, 53), androgen-targeting agents may not target all androgen-independent tumor cells, which could also harbor these methylation markers. Furthermore, most well-studied methylation markers were derived from earlier stages of PCa, whereas further epigenomic instability is known to occur with the mCRPC state (54).

Through sequential analysis using a genome-wide approach, we highlighted progressive changes in genes involved in tumorigenic processes in PCa and other cancers. Commonly altered genes in the promoter region included *PYCR1* and *SHARPIN,* which are known to be upregulated in PCa tissue and involved in processes such as proliferation and angiogenesis (55). Gene body methylation changes were observed in genes that regulate various tumorigenic processes, including *NFATC1* (47)*, PRDM16* (56), and *OPCML* (57). Pathway analysis also further highlighted key mechanisms that have been established in prostate tumor tissue studies, including the wnt pathway (16). However, the methylation patterns amongst these genes demonstrated extensive variability across patients and visits.

In order to further delineate key regions, we further assessed common regions that changed during treatment. These changes appeared to reflect dynamics of treatment response. For instance, patients that demonstrated CpG changes (up or down) in visit B and sustained the changes by visit C appeared to have a longer TTP. In contrast, there was a trend towards shorter TTP amongst patients that did not demonstrate sustained methylation changes. Among the genes included was Neuropeptides B/W receptor 1 (*NPBWR1*), which was shown to be upregulated in basal cells from prostate tissue (58), thus highlighting that genome-wide cfDNA methylation analysis is able to capture tumor dynamics and may further have additive potential to mutation analysis.

We also observed that many of the differentially methylated genes overlapped between Enzalutamide- and Abiraterone-treated patients. Although there are ongoing studies to compare efficacy of either treatment (59), the current consensus is that both perform similarly, and deciding between treatments in a patient-specific manner remains an ongoing challenge. We identified sets of genes altered in the promoter and gene body locations, including one large set amongst Enzalutamide treated patients. We were limited by the number of Abiraterone-treated patients, which may have limited heterogeneity, as seen in the larger Enzalutamide cohort. Further studies are needed to examine if the methylome is different between these treatment types. However, similar pathways were observed between the treatments, with neuronal pathways being the most prominent.

Given these neuronal-related genes/pathways, we wanted to examine the utility of these methylation changes in helping to predict treatment outcome, especially in identifying Neuroendocrine-CRPC. With our unique study design of sequential sampling, we could detect these methylation trends at earlier time points, demonstrating that the proportion of Neuroendocrine-like DMRs correlated with faster/shorter time to progression. In particular, the higher the proportion of certain DMRs with these NE-CRPC associated methylation patterns, the faster the time to progression. While we did not have confirmatory biopsies or other markers of NE disease (i.e. chromogranin A), increased methylation at Visit A could be utilized as predictive markers of response to androgen-targeting agents. Although we are currently limited to a single cohort, we now have a set of candidate regions that could be validated in additional cohorts.

## Conclusions

Our genome-wide sequencing protocol and analysis strategy has demonstrated the utility of intra-patient monitoring of cfDNA methylation. If changes in the methylome during treatment were sustained until clinical progression, these patients appeared to have a better prognosis with androgen-targeting treatment. Interestingly, methylation patterns associated with a highly aggressive form of the disease may serve as potential predictive markers. Further independent validation of these markers is needed to assess this predictive potential.

### Future Perspectives

Liquid biopsy based biomarkers will be playing a major role in how we treat patients with highly aggressive prostate cancer. Many studies focused primarily on genomic alterations associated with treatment resistance, but epigenomic markers could further explain the molecular heterogeneous mechanisms that contribute to disease progression. Ultimately, we anticipate that a combination of genomic and epigenomic markers will help facilitate optimal therapy decisions.

### Summary points

- We performed genome-wide methylation analysis of sequentially collected cfDNA samples from mCRPC patients receiving androgen-targeting therapy
- The DMRHunter pipeline was developed to identify differentially methylation regions between study visits for each patient
- Changes in promoter and gene body methylation was identified in key novel genes as well as previously established genes associated with tumourigenesis and the highly aggressive form of the disease, Neuroendocrine-CRPC
- Patients that were able to sustain methylation changes throughout treatment tended to present a longer time to clinical progression, suggesting that fluctuations in these CpG sites could help monitor response to treatment
- Elevated levels of Neuroendocrine-CRPC related methylation signals could serve potential predictive markers of resistance to androgen
- These findings could help with improving the clinical management of mCRPC using a minimally invasive approach

## Authors’ contributions

B.B. designed the study for intra-patient comparison. A.M.J. designed the clinical study for patient recruitment and sample collection. P.H. provided sample size calculations required for recruitment. M.R.P. performed sample processing, cfMeDIP-seq library preparation, and data analysis. M.B. developed and executed the DMRHunter pipeline and along with M.H., assisted with analysis to interpret cfMeDIP-seq datasets. R.I. and G.D.B. performed pathway analysis. S.Y.S. and D.D.D.C. designed the cfMeDIP–seq protocol and assisted with optimization for this study design. A.M.J., N.E.F. and A.R.H. were involved in patient follow-up and interpretation of data in the context of clinicopathological parameters. M.R.P., M.B. and B.B. composed the manuscript with feedback from all co-authors involved.

## Acknowledgements

We acknowledge the UHN Genitourinary clinic and Genitourinary Biobank for recruiting patients, collecting blood samples and for patient follow-up/data collection. All sequencing and initial data pre-processing was performed by the Princess Margaret Genomics Centre and the UHN Bioinformatics and HPC Core, Princess Margaret Cancer Centre, respectively. Differential methylation analysis was performed using the facilities provided by the Canada’s Michael Smith Genome Sciences Centre (British Columbia Cancer Agency). We also acknowledge Shivani Kamdar (SHS/LTRI) for assisting with sample processing and for helping to edit this manuscript.

## Ethics approval and consent to participate

All patients in this study provided informed written consent in accordance with approved institutional Research Ethics Board protocols from University Health Network (UHN) and Sinai Health System (SHS). Progress updates were submitted and approved annually by respective Research Ethics Boards. The study was performed in accordance with the Declaration of Helsinki. No personal information for any patient is presented in this manuscript. According to REB approvals, each patient was assigned unique study identifiers, which are not linked to personal health information.

## Availability of data and materials

All FASTQ and QC files will be made available in a public repository upon publication of this manuscript. Additional supplementary data files have also been included.

