## Supplementary Figures for "Dynamics of the cell-free DNA methylome of metastatic prostate cancer during androgen-targeting treatment"

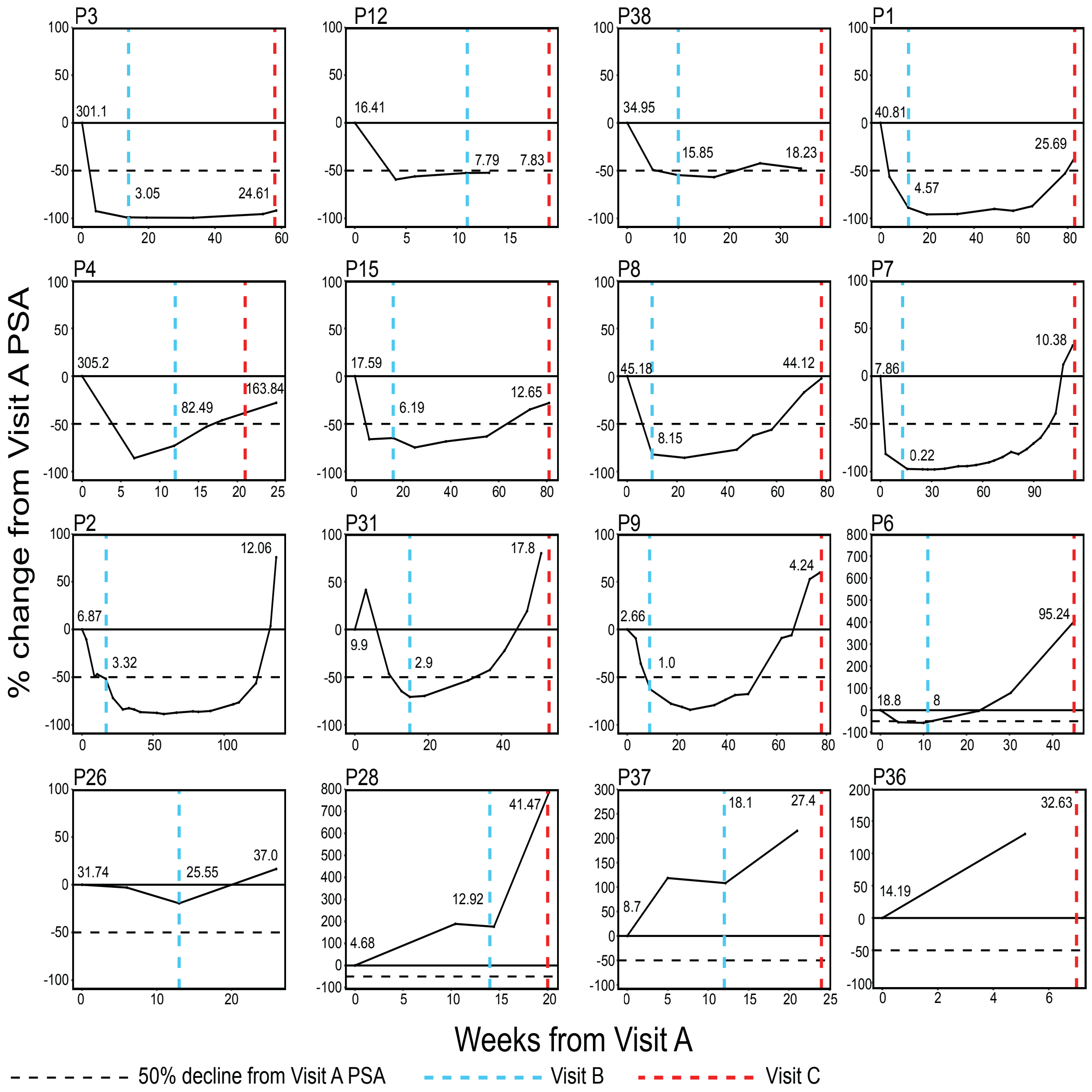


**Supplementary Figure 1. PSA dynamics for each patient.**
PSA levels for each patient were measured prior to starting treatment and typically monthly following initiation of either Abiraterone or Enzalutamide. For each patient, the percent change in PSA value from baseline/Visit A is plotted against the number of weeks from Visit A. The horizontal black dashed line indicates favorable PSA response (50% decline from Visit A). The vertical blue and red lines mark when Visits B and C were collected, respectively. PSA values (ug/L) are shown for each visit.

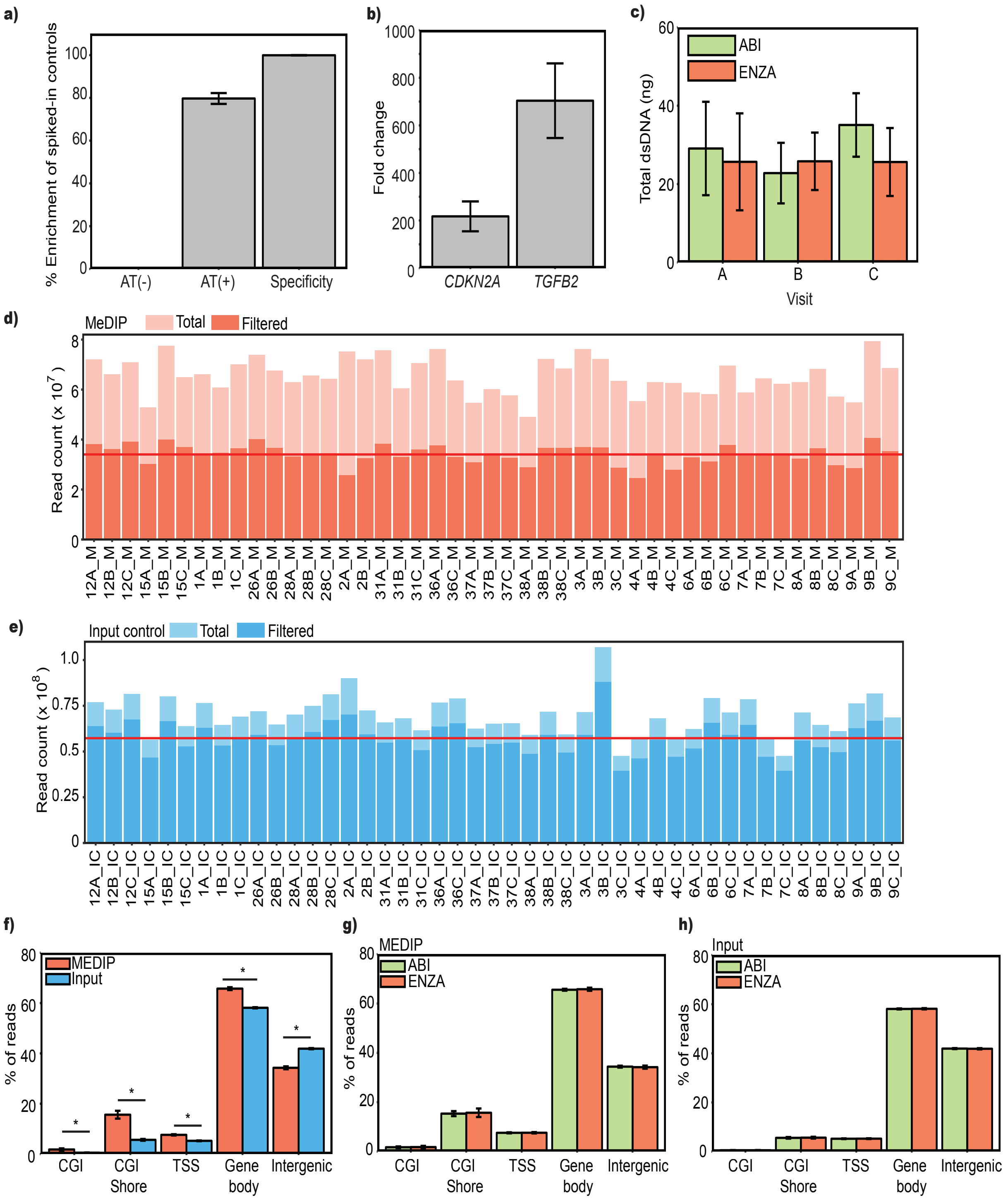


**Supplementary Figure 2. cfMeDIP-seq quality controls and filtering.**
The cfMeDIP-seq protocol was first tested on low amounts of 22Rv1 DNA (50 ng) sheared to similar size range as cfDNA. (a) Control methylated (+) and unmethylated (-) Arabidopsis thaliana (AT) DNA was spiked into each sample prior to immunoprecipitation and qPCR was

performed of MeDIP enriched and input control DNA. The proportion of unmethylated, methylated and specificity (methylated:unmethylated AT DNA ratio) is shown (N=3). (b) qPCR assessment of known methylated genes in 22Rv1 (*CDKN2A* and *TFGB2*) are expressed as fold change over a known unmethylated gene (*HOXD8*) (N=3). (c) The average amount of double stranded cfDNA is summarized across all visits collected (45 samples total) and further stratified by treatment type (standard deviation is shown). MeDIP-seq was performed on 45 MeDIP libraries and 45 corresponding input control libraries. All reads were mapped to the reference genome, duplicates collapsed and low quality reads removed. The total versus filtered read counts is shown for each patient and visit in (d) MeDIP libraries and (e) input control libraries, with average read count shown (red line). (f) The average proportion of MeDIP and input control reads are shown across CpG islands (CGI), CGI shores, transcriptional start sites/promoters (TSS), gene bodies and intergenic regions. Significant differences among MeDIP vs input read proportions are shown using the Mann-Whitney U-test ($P<0.05$). Proportion of reads across all regions comparing Enzalutamide and Abiraterone groups are shown for (**g**) MeDIP and (**h**) input controls.

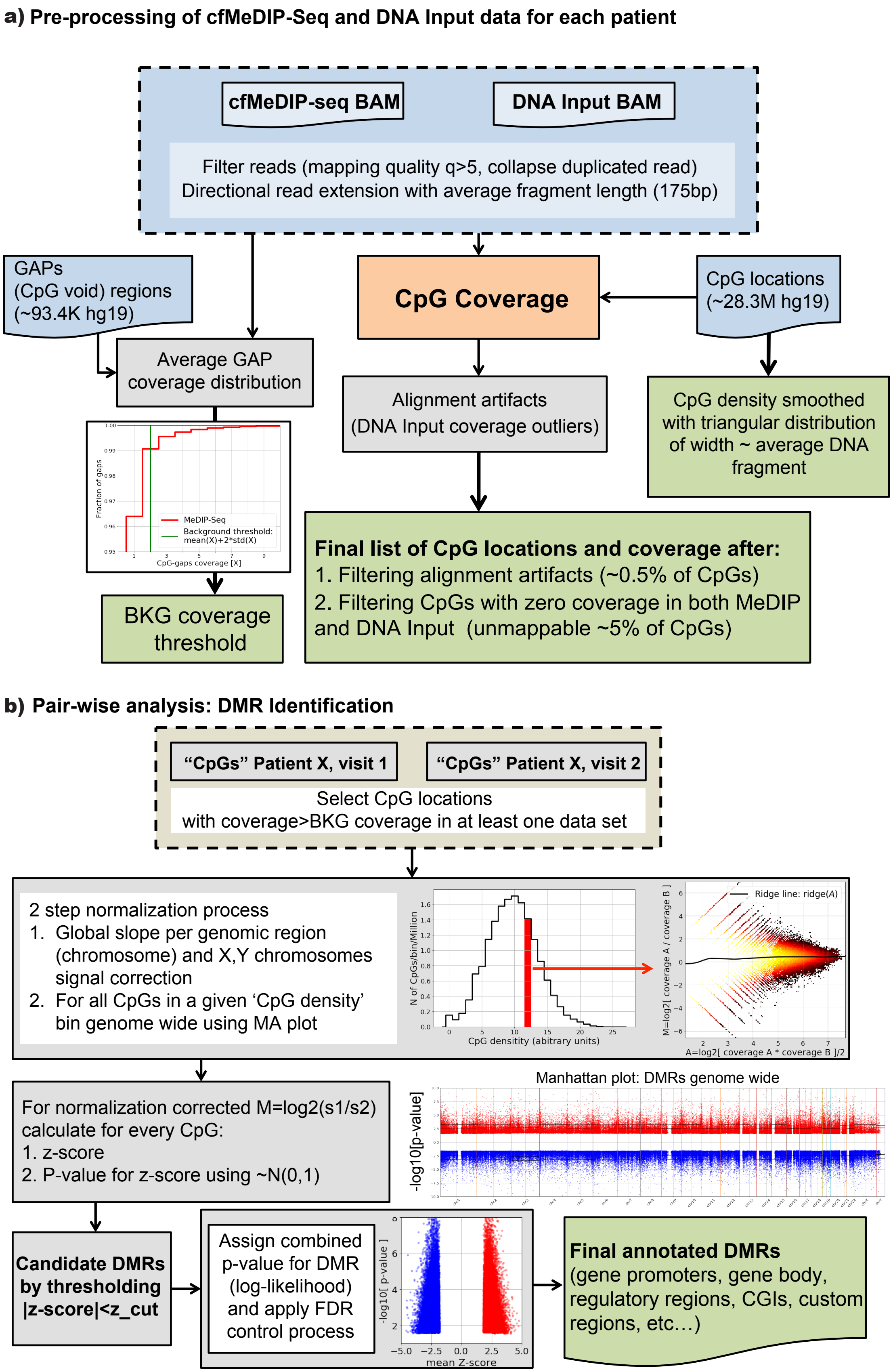
a) Pre-processing of cfMeDIP-Seq and DNA Input data for each patient
cfMeDIP-seq BAM
DNA Input BAM
Filter reads (mapping quality q>5, collapse duplicated read)
Directional read extension with average fragment length (175bp)
GAPs
(CpG void) regions
(~93.4K hg19)
CpG Coverage
CpG locations
(~28.3M hg19)
Average GAP
coverage distribution
Fraction of gaps
MeDIP-Seq
Background threshold:
mean(X)+2*std(X)
CpG-gaps coverage [X]
BKG coverage
threshold
Alignment artifacts
(DNA Input coverage outliers)
CpG density smoothed
with triangular distribution
of width ~ average DNA
fragment
Final list of CpG locations and coverage after:
1. Filtering alignment artifacts (~0.5% of CpGs)
2. Filtering CpGs with zero coverage in both MeDIP
and DNA Input (unmappable ~5% of CpGs)
b) Pair-wise analysis: DMR Identification
“CpGs” Patient X, visit 1
“CpGs” Patient X, visit 2
Select CpG locations
with coverage>BKG coverage in at least one data set
2 step normalization process
1. Global slope per genomic region
(chromosome) and X,Y chromosomes
signal correction
2. For all CpGs in a given ‘CpG density’
bin genome wide using MA plot
N of CpGs/bin/Million
CpG densitity (abitrary units)
Ridge line: ridge(A)
M=log2[ coverage A / coverage B ]
A=log2[ coverage A * coverage B ]/2
For normalization corrected M=log2(s1/s2)
calculate for every CpG:
1. z-score
2. P-value for z-score using ~N(0,1)
Manhattan plot: DMRs genome wide
-log10[p-value]
Candidate DMRs
by thresholding
|z-score|<z_cut
Assign combined
p-value for DMR
(log-likelihood)
and apply FDR
control process
-log10[ p-value ]
mean Z-score
Final annotated DMRs
(gene promoters, gene body,
regulatory regions, CGIs, custom
regions, etc…)

**Supplementary Figure 3. Summary of cfDNA DMR hunter pipeline.**
(a) Prior to initiating DMR analysis, stringent read quality filtering was applied and reads were extended. Coverage of all CpG sites in the genome was calculated for MeDIP libraries and removal of regions without coverage (using Input control libraries). (b) Pairwise analysis was performed to identify differentially methylated CpG sites followed by identification of DMRs and annotation. More detailed description of cfDNA DNR hunter in supplementary methods.

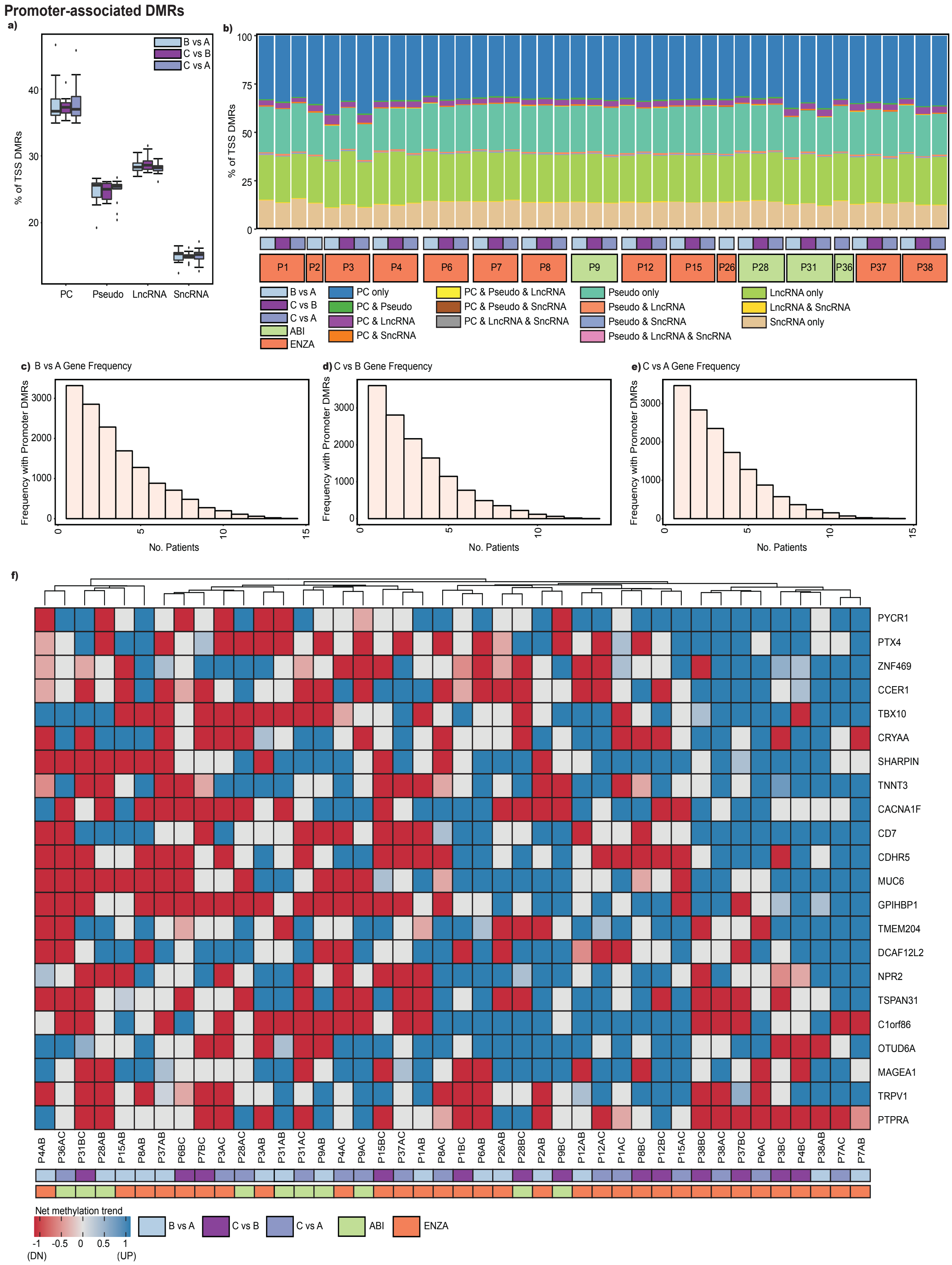


**Supplementary Figure 4. Distribution of TSS/promoter-associated DMRs in all patients.**

(a) The proportion of TSS/promoter-associated DMRs near protein coding (PC), pseudogenes (Pseudo), long non-coding RNA (lncRNA) and short non-coding RNA (sncRNA) was calculated for all visit comparisons. The median, first and third quartile is shown. (b) For each patient, the relative proportion of all TSS-associated DMRs for each genomic region or combination is shown. The frequency/recurrence of protein coding promoters with DMRs across all patients in (c) B vs A, (d) C vs B and (e) C vs A. (f) The net methylation change (ratio of increased to decreased methylation) for all comparisons was calculated and heatmap shows overall methylation changes for promoter regions that were altered in 30 or more visit comparisons.

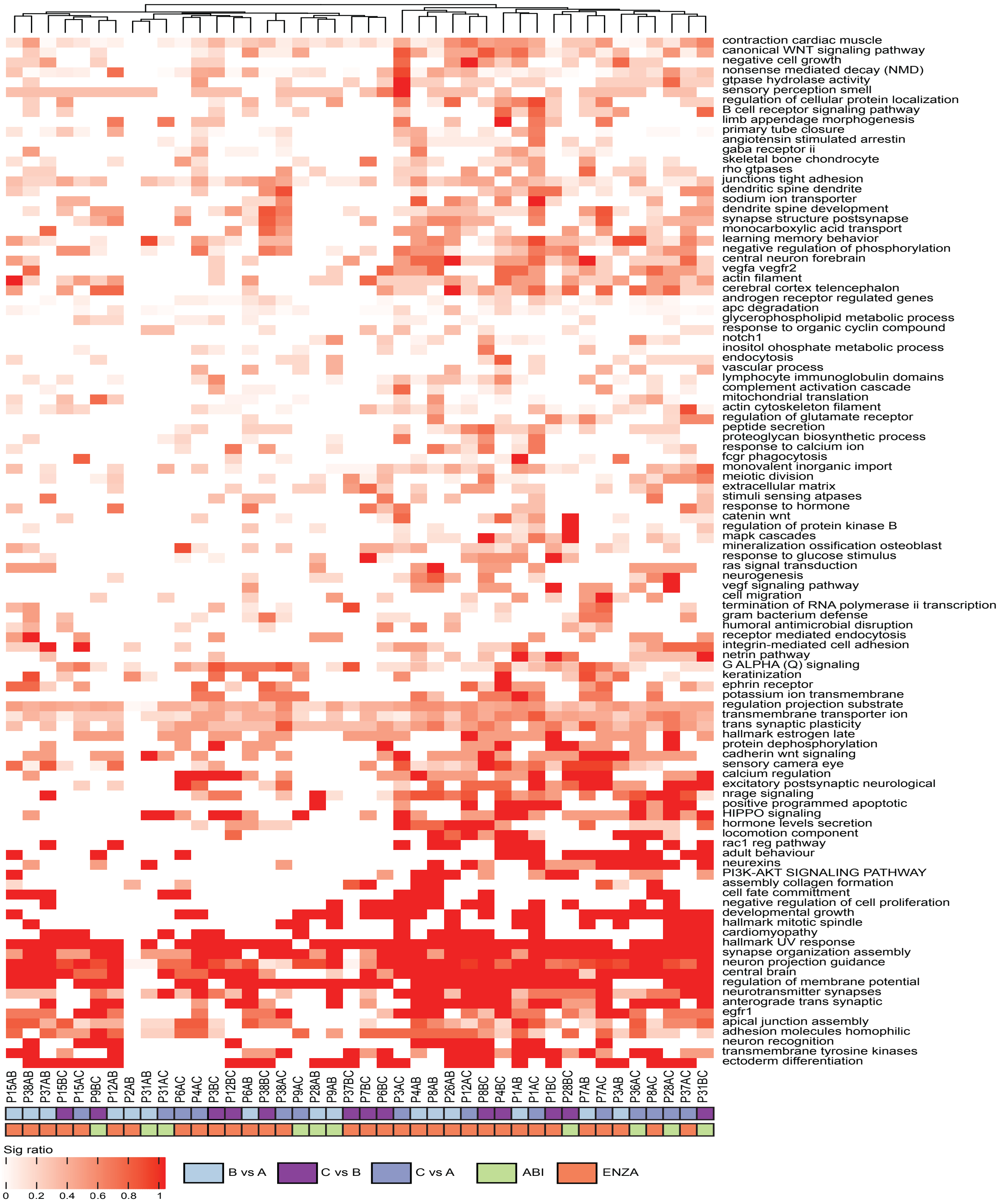


**Supplementary Figure 5. Pathway analysis of promoter regions with increased methylation during treatment.**

Pathway analysis was performed for promoter regions with increased methylation (UP) during treatment for all patients and comparisons. Common pathways were grouped together into themes, and the ratio of the number of significant pathways to the total number of pathways in each theme was calculated and represented in this heatmap. The brighter the color, the more significant pathways within the specific theme (labeled on the right).

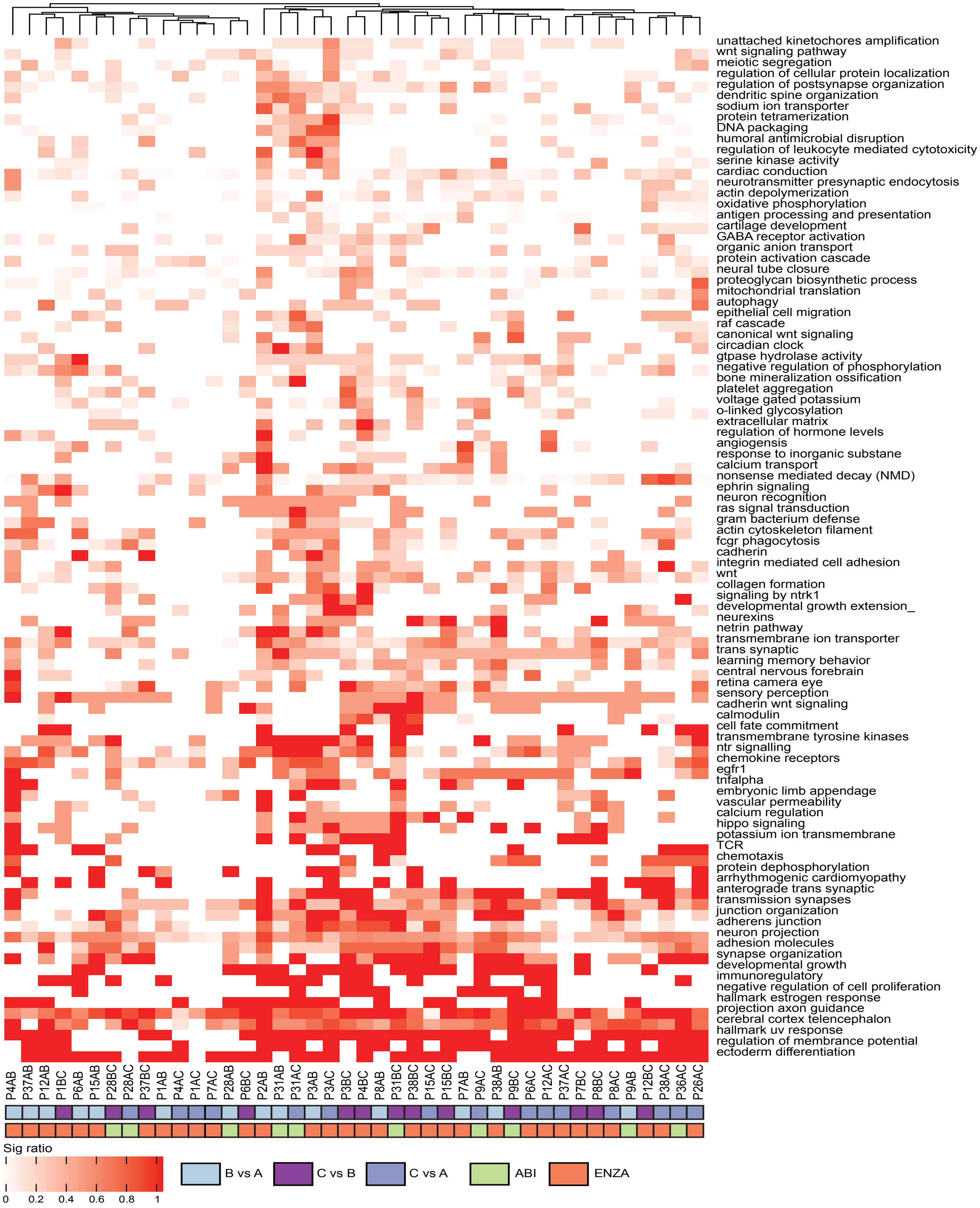


**Supplementary Figure 6. Pathway analysis of promoter regions with decreased methylation during treatment.**

Pathway analysis was performed for promoter regions with decreased methylation (DOWN) during treatment for all patients and comparisons. Common pathways were grouped together into themes, and the ratio of the number of significant pathways to the total number of pathways in each theme was calculated and represented in this heatmap. The brighter the color, the more significant pathways within the theme (labels on the right).

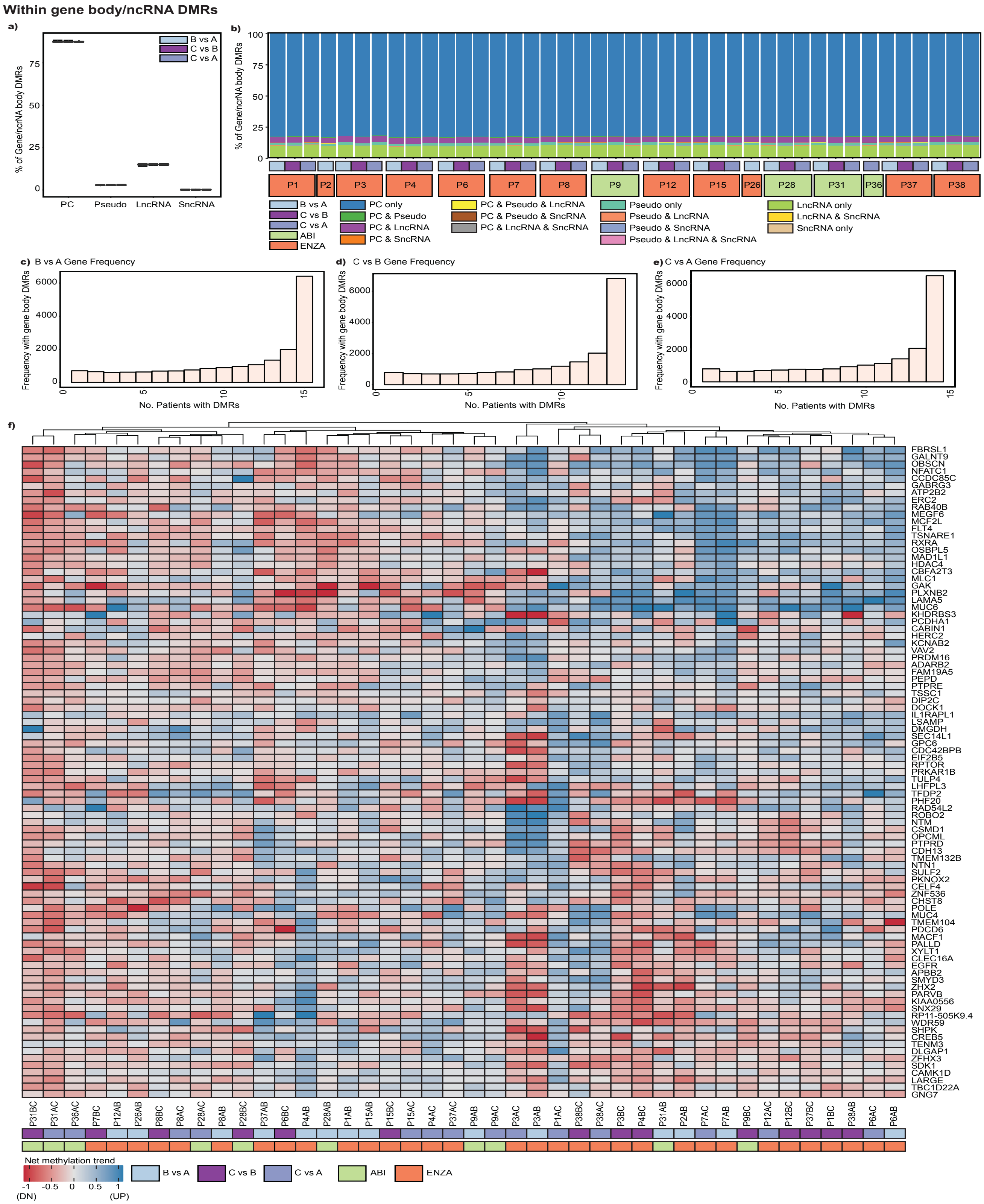


**Supplementary Figure 7. Distribution of DMRS within gene bodies and ncRNA.**

(a) The proportion of DMRs within protein coding (PC) genes, pseudogenes (Pseudo), long non-coding RNA (lncRNA) and short non-coding RNA (lncRNA) was calculated for all visit comparisons. The median, first and third quartile are shown. (b) For each patient the distribution of all DMRs for each genomic region or combination is shown. The frequency of protein coding gene bodies with DMRs in (c) B vs A, (d) C vs B and (e) C vs A. (f) Heatmap shows overall methylation changes within protein coding regions that were altered in all visit comparisons.

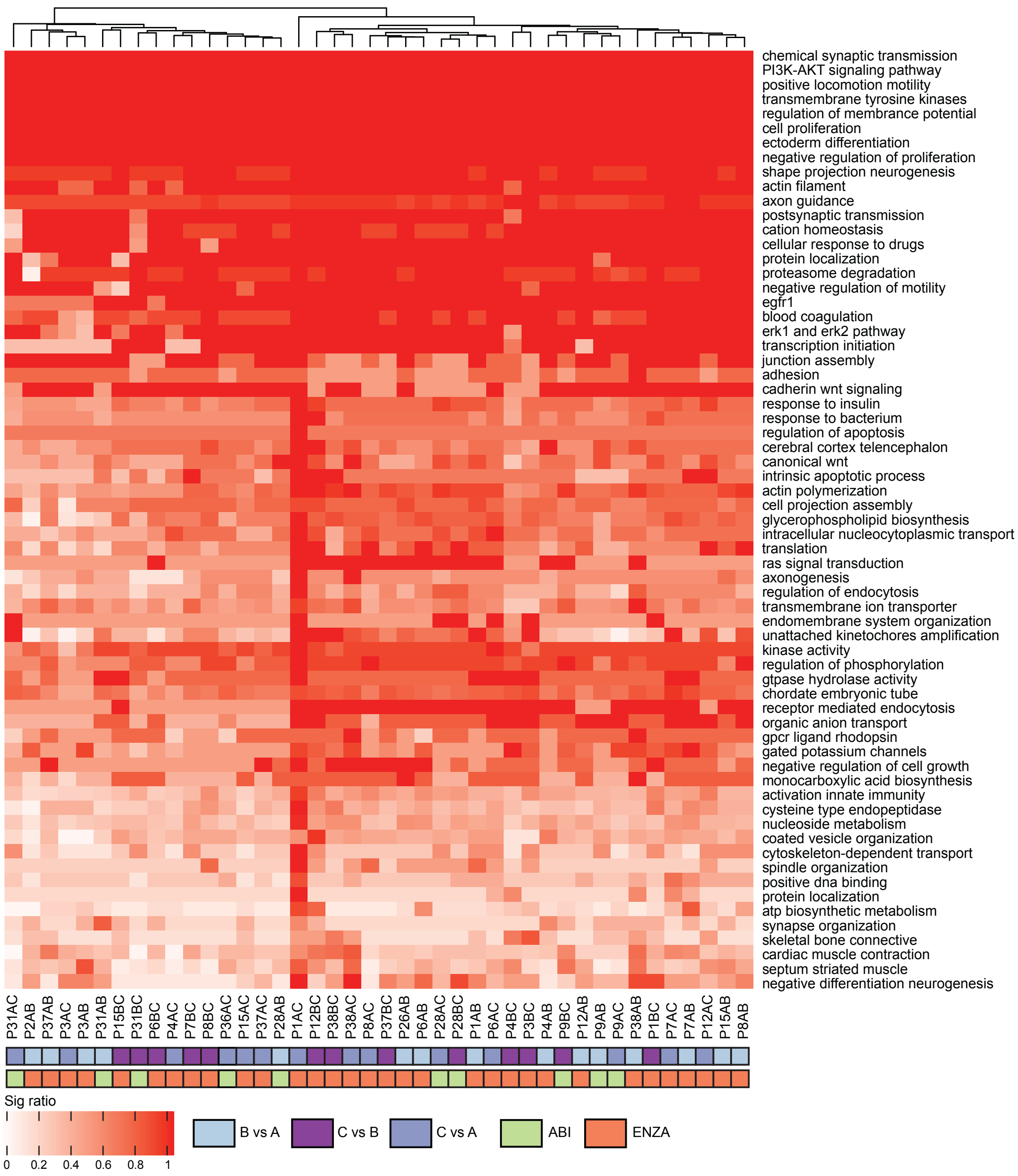


**Supplementary Figure 8. Pathway analysis of gene body regions with increased methylation during treatment.**

Pathway analysis was performed for each patient/comparison and for gene body regions that demonstrated increased methylation during treatment. Common pathways were grouped together into themes. For each theme, the ratio of the number of significant pathways to the total number of pathways in the theme was calculated and represented in this heatmap.

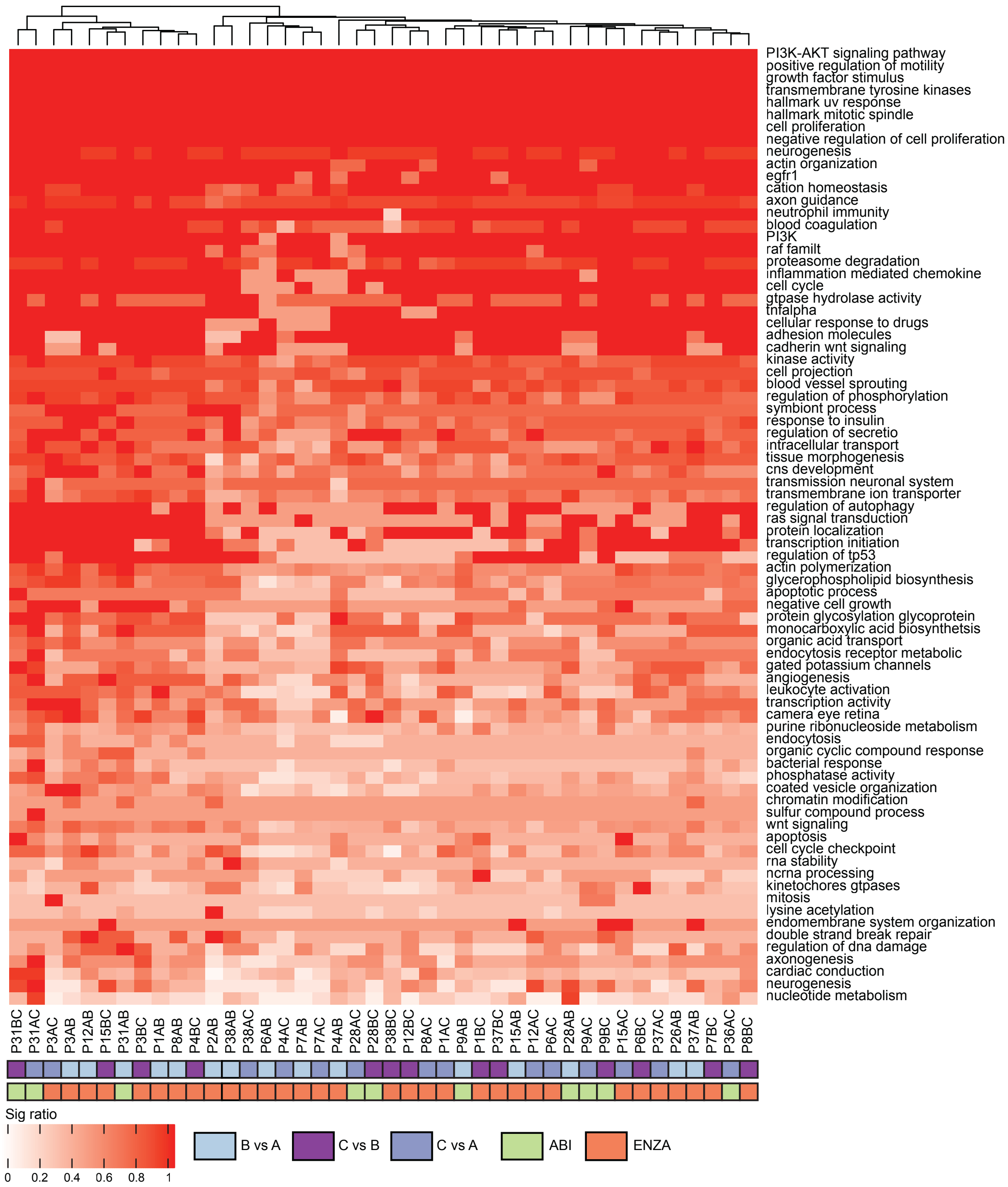


**Supplementary Figure 9. Pathway analysis of gene body regions with decreased methylation during treatment.**

Pathway analysis was performed for each patient/comparison and for gene body regions that demonstrated decreased methylation during treatment. Common pathways were grouped together into themes. For each theme, the ratio of the number of significant pathways to the total number of pathways in the theme was calculated and represented in this heatmap.

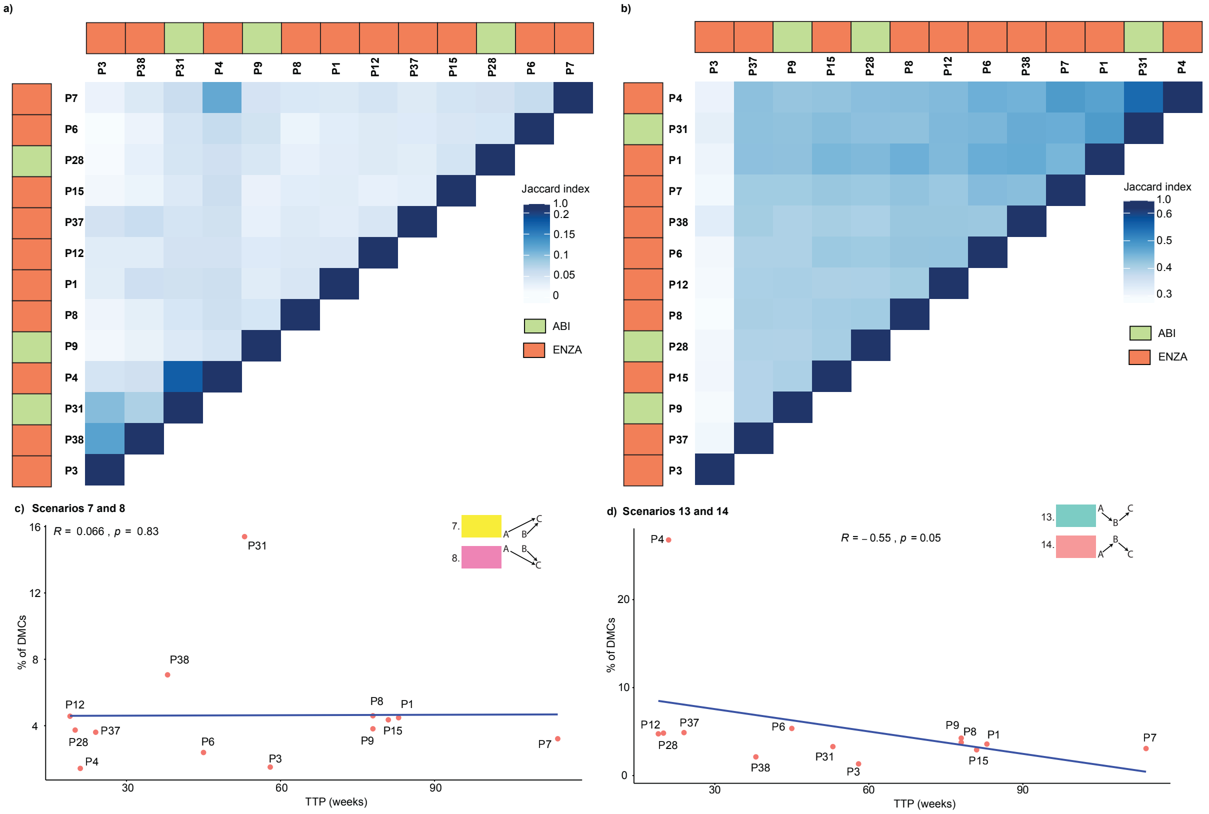


**Supplementary Figure 10. Distribution of genes with shared CpG sites.**
(a) For promoter sites, genes associated with CpG sites that were altered in two or more visits were compiled for each patient with 3 visits. Heatmap shows Jaccard index coefficient assessing the degree of similarity amongst all patients. The more genes shared by patients, the darker the shade of blue (b) Similarly, Jaccard index coefficient analysis was performed for gene body regions. Spearman correlation analysis was perform to assess relationship of TTP with (c) proportion of DMCs with increased/decreased methylation in C vs A/B or (d) DMCs that increased/decreased in B vs A and returned to similar levels in C as A.
