## Supplementary Methods for "Dynamics of the cell-free DNA methylome of metastatic prostate cancer during androgen-targeting treatment"

### Supplementary Methods Section – cfDNA DMRHunter and Pathway Analysis

#### cfDNA DMRHunter

DMRHunter was designed to perform a pair-wise comparison of two MeDIP-seq data sets to find significantly differentially methylated regions (**DMRs**) from continuous blocks of differentially methylated CpGs (**DMCs**), as well as to annotate DMRs with various overlapping genomic features (e.g. genes, CpG Islands). DMRHunter uses MeDIP-seq and DNA-Input control aligned reads in the BAM format files. The only significant parameter is a user defined FDR threshold. Analysis consists of a multiple steps described below and summarized in **Supplementary Figure 3**.

#### 1. CpG coverage calculation and CpG filtering

Using the hg19 reference genome sequence, we first generated a resource file with positions of all CpGs (28,245,162 CpGs on assembled chromosomes). For all studied MeDIP-seq and DNA Input control data, we calculated total coverage for each CpG site. For single-end sequencing data (which was the case in this study), each aligned read was directionally extended by the average DNA fragment length (175 bp for cfDNA) and coverage of extended reads was evaluated at CpG locations using in-house custom tools BAM2WIG and RegionsCoverageFromWIGCalculator (see GSCtools at http://www.epigenomes.ca/tools-and-software). Note: prior to coverage calculation low quality reads (BWA QA<5) were filtered/removed and duplicated reads were collapsed – only one copy was considered. For pair-end data, DMRHunter uses aligned DNA fragments for coverage calculation.

We then excluded CpGs within potentially unmappable genomic regions (likely repeats). That is, CpGs with zero coverage in both MeDIP-seq and DNA Input data for both samples used in the pair-wise comparison. Typically, ~5% of CpGs were flagged as 'unmapped' and excluded in each comparison.

In addition, we excluded CpGs that have extremely high coverage in both DNA Input data sets (selecting those that have coverage > 99.5 percentile). Those are potentially CpGs within regions of "alignment artifacts" (regions showing spurious enrichment in most of NextGen sequencing data) (1). This accounted for <1% of CpGs that were also excluded. It is important to filter CpGs potentially falling into unmappable and 'alignment artifact' regions at the earliest stage of the analysis, as keeping them would affect all downstream calculations.

#### 2. Finding threshold for MeDIP-seq coverage to exclude noise

In order to separate background noise from a true methylation signal in MeDIP-seq data, we defined 'CpG void' regions that were at least 500bp distance apart with closest CpG site, which we referred to as 'gap' regions. We found 93,419 such 'gap' regions genome wide, with total length 260,299,662 bp.

As our estimated cfDNA fragment length is ~200bp, and assuming that methylation happens predominately in CpG context, experimentally observed coverage of the 'gap' regions is mostly due to the background noise.

This was used to estimate background coverage threshold values, which we defined as:

*bkg_threshold = mean[gap coverage]+2*std[gap coverage].*

This threshold was determined individually for each sample. After applying *bkg_threshold* to the CpG coverage, we found that typically ~40% of CpGs did not have detectable MeDIP signal in both samples in our data.

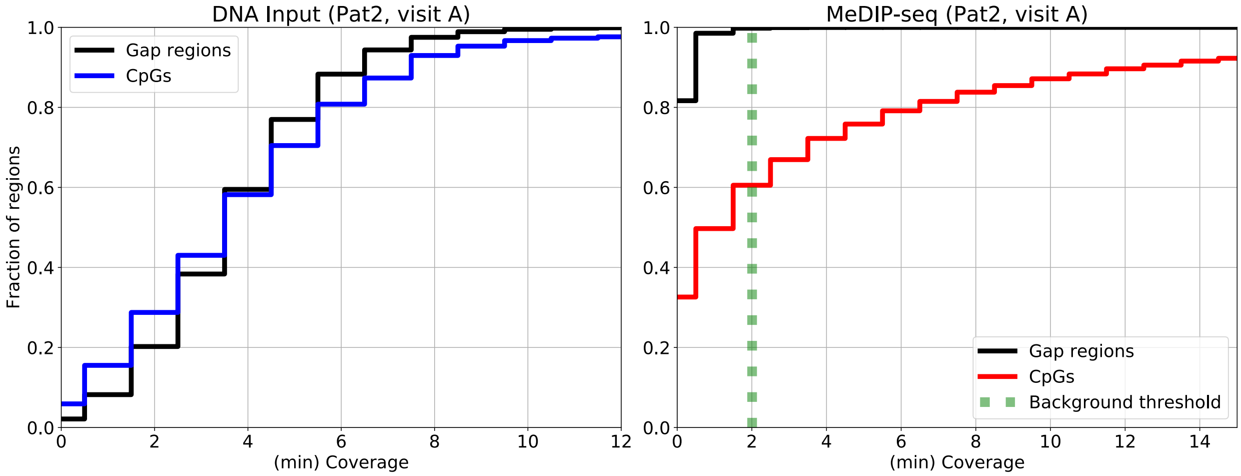


**Fig SM1.** Example of coverage distributions for MeDIP-seq and DNA Input for P2 visit A. Left panel shows mean coverage of 'gap' regions and CpGs in DNA Input data, while right panel shows MeDIP-seq data, as well as evaluated background threshold. As expected, the DNA Input coverage at CpGs is similar to gap regions.

### <u>3. Estimation of CpG density</u>

Distribution of CpG locations in human genome is highly variable, some CpGs are very sparse others in dense clusters (e.g. CpG Islands,CGIs) (2) (See below).

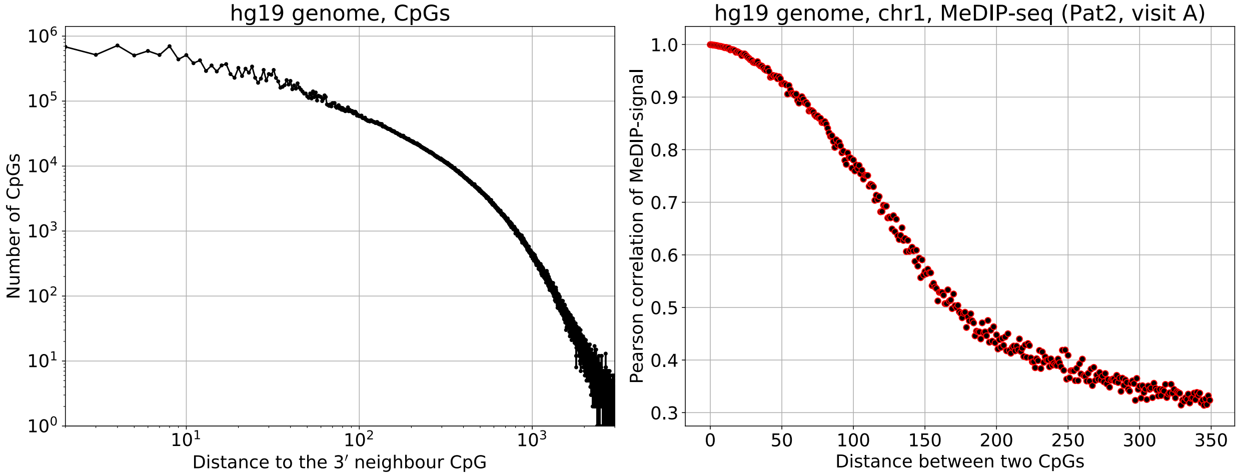


**Fig SM2.** The distribution of distances between 2 neighboring CpGs is shown across the genome (left panel). ~79% of CpGs have at least one neighboring CpG site at a distance less than 150bp, while ~3.4% of CpGs have the closest neighbor >500bp away. The fragment length of precipitated DNA limits resolution of a MeDIP-seq experiment, resulting in high correlation between coverage of CpGs with relative distance less than DNA fragment. The right panel shows the Pearson correlation for CpGs MeDIP-seq coverage (on chr1) as a function of a distance between CpGs for P2 visit A. Correlation was calculated for a set of genomic CpGs for which another CpG at a given distance was found. As expected, the correlation is high for proximal CpGs and drops rapidly at ~175bp (approximately precipitated DNA fragment length).

To control variability in CpGs positions, we assigned 'CpG density' value for every CpG. We smoothed an array of zeros and ones (with 1 being placed at CpG coordinates genome wide) using a triangular kernel with a mean DNA fragment as a half-width parameter. 'CpG density' was defined as value of the smoothed array at the CpG location.

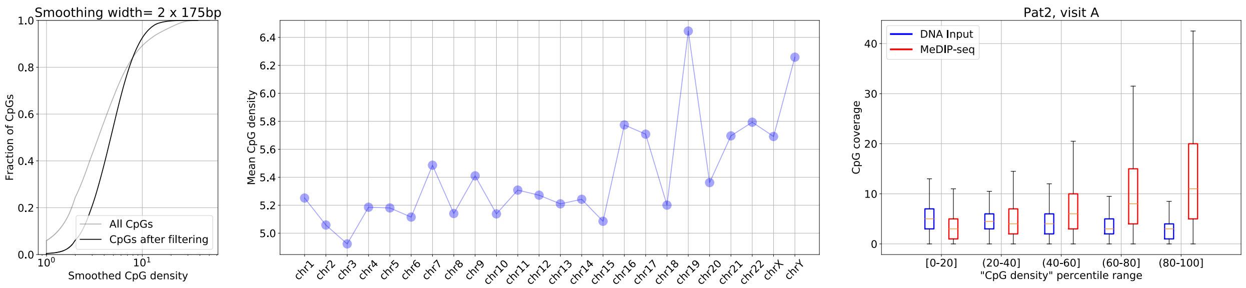


**Figure SM3.** Left panel shows cumulative distribution of the 'CpG density', which with our normalization ranges from one for an isolated CpG, up to ~51 for CpGs in the dense clusters. The middle panel shows mean 'CpG density' for each chromosome, with gene-rich chr19 having highest CpG density. We observed that properties of MeDIP-seq data vary depending on 'CpGs density' as illustrated in the right panel. Coverage distributions for both DNA Input and MeDIP-seq are shown for CpGs from 5 different percentile intervals of 'CpG density'. While, as expected, coverage distribution for DNA Input is invariant, MeDIP-seq coverage depends strongly on the 'CpG density' range, with both mean and variance of coverage increasing with increasing 'CpG density'.

In the next step of analysis, we decided to group CpGs with similar 'CpG density' for normalization between two data sets,

### 4. CpG coverage normalization for two samples

Proper normalization is a key for MeDIP-seq sample comparisons and determination of differentially methylated regions (DMRs). Two compared libraries can typically have different sequencing depth as well different biases (originating from all experimental steps, including IP, library construction and sequencing). We aimed to control some of these biases by sequencing samples for different visits from the same patient together. We observed (from all MeDIP-seq samples analyzed) that not only the absolute value of CpG coverage depends on 'CpG density' (as discussed above, **Figure SM3**), but also relative normalization between two samples depends on the CpG density – see left panel on Fig SM4 that shows distributions of log2-transformed ratio of the CpG coverage for visits A and B for patient 2. Our normalization strategy is based on the assumption that methylome is relatively similar for two data sets and that majority of CpGs are not differentially methylated.

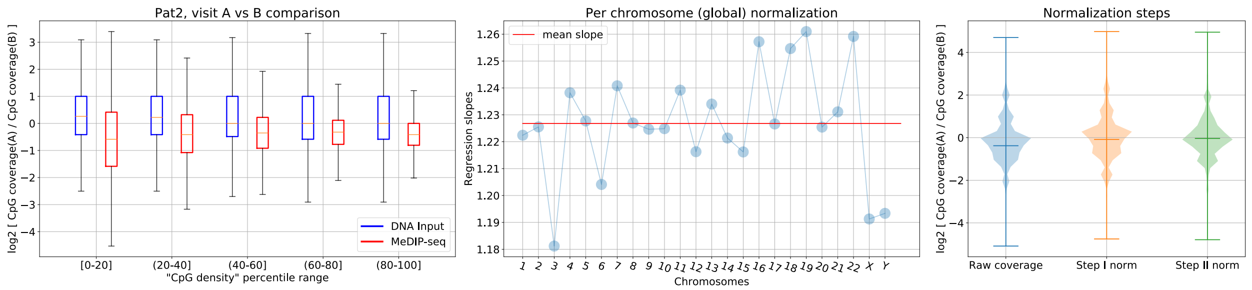


**Figure SM4.** Illustration of normalization procedure**.** Left panel shows distributions of log2-transformed ratio of the CpG coverage for visits A and B for patient 2 calculated 5 different percentile intervals of 'CpG density'. Plot clearly shows 'CpG density' dependence of coverage ratio for MeDIP-seq signal. Middle panel showing regression slopes for each chromosome used in the first normalization step. We observed some variability between different chromosomes.
Right panel show violin plots for the log2-ratio of the CpG coverage between two visits before the normalization and after two normalization steps.

The two-step normalization procedure was as follows:

1. We first corrected CpG coverage on sex chromosomes for male samples by multiplying measured values by 2 two to compensate for a single copy of X and Y chromosomes relative to autosomes and performed global per genomic region normalization (in our case we used entire chromosome). In this step we take into account the difference in total number of reads in each region between two samples. Technically we performed a linear regression of MeDIP-seq coverage at CpGs between two samples (see Fig SM4 middle panel) and used a regression slope to introduce relative normalization between samples. The choice of genomic regions could be different, e.g. one can use regions of distinct copy-number values for samples with strong copy-number variability.
2. In the next step, we split CpG density into 100 bins (using percentile) and re-normalized data in each density bin. To do this, we considered MA plot (**See Supplementary Figure 3**) and assumed that majority of CpGs in each bin are not differentially methylated and should have the same normalized CpG coverage. M and A are defined for every CpG_i, using MeDIP-seq coverage values for two samples:

   $$A(CpG\_i)=log2(coverage1*coverage2)/2,$$
   $$M(CpG\_i)=log2(coverage1coverage2),$$

   We fit positions of the ridge in the MA plot, ***M_ridge=F(A),*** to infer the normalization coefficient for every value of A.

To validate the normalization process we demonstrated that CpG coverage distributions genome wide in both samples are very similar (Jensen-Shannon distance <0.01) (3). Right panel on Fig SM4 shows final results of normalization process. One can see this example that after the second normalization step median of the distribution became very close to zero and the shape of the distribution is more symmetrical around zero.

At this step we also assign a Z-score for the M-variable:

$$Z(A)=(M(A)-M\_ridge(A))/std(M(A)),$$

and using a normal distribution $\sim N(0,1)$ we assigned p-values for a Z-score for every CpG.

**5. Defining DMRs**

We used the Z-score threshold:

$$|Z\text{-}score|>1.96,$$

to first select 'islands' of DMCs (continuous blocks of DMCs) that are forming potential DMR candidates.

Using p-value for every DMC in DMR candidates, we calculate a combined p-value while taking into account correlation between proximal CpGs using a previously developed methodology (4). By calculating combined p-value we make it possible to apply further FDR control process to DMRs of different sizes, ranging from a single CpG to hundreds of blocked CpGs. Note that this approach does not require *ad-hoc* parameters, such as DMC proximity threshold or minimal number of CpGs per DMR, which are typically used in many DMR defining software tools.

For the False Discovery Control we used Benjamini-Hochberg procedure with FDR=0.01 (5). For the final set of DMRs, DMRHunter also outputs all DMCs falling into DMRs.

**6. Annotation**

Final step is annotating DMRs with any list of genomic features (genome feature contains id/name and genomic coordinates). Every DMR is annotated with a genomic feature if it overlaps genomic feature by at least one base, such as CGIs and genes (6).

DMRHunter tool is a stand-alone program written in Python 3.5 and typical execution time for a single pair-wise comparison on a single CPU is ~30-60mins.

**Pathway Analysis**

To define differentially regulated pathways in our different sets of DMRs. For each patient there were three unique sets of DMRs (one for each of the comparisons between visits B vs A, C vs A and C vs B) which were analyzed for enriched pathways. A local implementation of the Genomic Regions Enrichment of Annotations Tool (GREAT) (7) algorithm was used to calculate the set of enriched pathways using an updated pathway definition file from October 01, 2018 (http://download.baderlab.org/EM_Genesets/). The pathways were obtained from MsigDB-c2, NCI, Biocarta, IOB, Netpath, HumanCyc, Reactome, Panther and Gene Ontology (GO) databases.

The local implementation of GREAT performed both hypergeometric and two-sided binomial test using the foreground and background set of DMRs, where the foreground included only regions that were significantly differential and the background contained all regions found in both visit for the given patient (i.e. for P1 B vs A comparison the background consisted of all regions found in P1 for visit B vs visit A). Genomic coordinates were downloaded from Ensembl and were resized to include 1500 bp upstream from the gene definitions. P-values were further corrected using Benjamini-Hochberg.

For each patient and each comparison a set of enriched pathways are computed. A pathway is considered enriched for a given patient if the corrected hypergeometric and corrected binomial p-value were both less than 0.05. Given the amount of pathways (n=42) associated with our set, we summarized to a set of themes using the AutoAnotate App (8) in in Cytoscape (v3.6.1) to help compare the results between different patients and visits. The AutoAnnotate App creates themes from highly connected pathways (shared genes) reducing the cluster to the most frequent words in the pathway names for the region. A patient and visit is considered to be part of a computed theme if it is enriched for any pathway associated with the theme. For each theme, the ratio of significant pathways to total pathways was calculated and represented as heatmaps.
